# The *vprAB-ompV-virK* operon of *Vibrio cholerae* senses antimicrobial peptides and activates the expression of multiple resistance systems

**DOI:** 10.1101/2024.08.28.609885

**Authors:** Annabelle Mathieu-Denoncourt, Gregory B. Whitfield, Antony T. Vincent, Julien Pauzé-Foixet, Feriel Mahieddine, Yves V. Brun, Marylise Duperthuy

## Abstract

Antimicrobial peptides are small cationic molecules produced by eukaryotic cells to combat infection, as well as by bacteria for niche competition. Polymyxin B (PmB), a cationic cyclic antimicrobial peptide, is used prophylactically in livestock for infection prevention and as a last-resort treatment for multidrug-resistant bacterial infections in humans. In this study, a transcriptomic analysis in *Vibrio cholerae* showed that expression of the uncharacterized gene *ompV* is stimulated in response to PmB. We found that *ompV* is organized in a conserved four-gene operon with the two-component system *vprAB* (*carRS*) and *virK* in *V. cholerae*, and that these genes are also upregulated in response to PmB treatment. A *virK* deletion mutant was more sensitive to the human cathelicidin LL-37 than the wild-type strain, while an *ompV* mutant was more sensitive to PmB and LL-37, suggesting that both OmpV and VirK contribute to antimicrobial resistance in *V. cholerae*. This increased sensitivity to antimicrobial peptides was not due to membrane destabilization or reduced sequestration by membrane vesicles as a result of *ompV* deletion. Instead, our transcriptomic analysis showed that the efflux pump *vexAB*, a known effector of PmB resistance, was also upregulated in the presence of PmB in an *ompV*-dependent manner. Examination of the predicted structure of OmpV revealed a lateral opening in the β-barrel wall with access to an electronegative pocket in the barrel lumen that can accommodate PmB. Such an interaction could facilitate intracellular signaling through a conformational change in OmpV. This is the first evidence of a specialized operon governing multiple systems for antimicrobial resistance in *V. cholerae*.

**Author Summary:** In this study, we identified the first specialized operon controlling multiple systems of antimicrobial resistance in *V. cholerae*. The operon encodes the two-component system *vprAB*, which activates the main mechanism of polymyxin B resistance in *V. cholerae*, and the uncharacterized genes *ompV* and *virK*. We provide evidence that OmpV and VirK are implicated in antimicrobial resistance and show that OmpV has a membrane-accessible lateral opening into a pocket that could accommodate the antimicrobial peptide polymyxin B. We propose that OmpV acts as an outer membrane sensor that signals the presence of antimicrobial peptides to activate the expression of the operon, leading to the activation of multiple mechanisms of resistance, including modifications of the outer membrane and the multi-drug efflux system *vexAB*.

## Introduction

Antimicrobial peptides (AMPs) are cationic molecules of low molecular weight with activity against bacteria, viruses, and fungi (1). They are produced by eukaryotes for immune regulation and to maintain the homeostasis of the microbiota, and by bacteria for competition for environmental niches (2–4). The ionic interaction of cationic AMPs with the negatively charged bacterial membrane leads to pore formation, leaking of intracellular content, and eventually death (5). AMPs can also have many intracellular targets. Polymyxin B (PmB) is a non-ribosomal cyclic antimicrobial peptide produced by *Paenibacillus polymyxa* that is used as a last resort treatment for multidrug resistant Gram-negative bacterial infections (6). Polymyxins are poorly absorbed during treatment and can thus be excreted and accumulate in the environment, which could lead to the development of resistance (7, 8).

*Vibrio cholerae* is a Gram-negative bacterium that resides in aquatic environments (9). It is responsible for the disease cholera, caused by consumption of contaminated water or food. *V. cholerae* is divided into 200 serogroups, of which only O1 and O139 cause cholera (10). The O1 serogroup is further divided into 2 biotypes, Classical, responsible for the first six cholera pandemics, and El Tor, which is responsible for the 7^th^ ongoing pandemic (10). The α-helical cathelicidin LL-37 and several α and β-defensins are produced by intestinal cells in response to *V. cholerae* infection (11). Unlike the Classical strains, El Tor strains can resist PmB by decreasing the negative charge of their outer membrane through *almEFG*, thus reducing interactions with cationic AMPs. AlmG is a glycyl transferase responsible for aminoacylation of the lipopolysaccharide (LPS) and represents the main mechanism of resistance to PmB in those strains (6). The expression of *almG* in response to PmB is regulated by the two-component system VprAB, also called CarRS, in which VprB is the sensor histidine kinase activated by periplasmic AMPs and VprA is the response regulator (12–15). Other mechanisms, such as efflux pumps, also contribute to AMP resistance in *V. cholerae* (16). The efflux pumps belonging to the resistance-nodulation-division (RND) transporter family are tripartite drug-ion antiporters spanning both the inner and outer membranes, facilitating the transport of substrates from the cytoplasmic membrane or periplasm to the extracellular milieu (17, 18). Six RND transporters are encoded in *V. cholerae* with different substrate specificity (19), of which VexAB is associated with resistance to detergents and antimicrobials such as PmB (16).

OmpV is a protein of 28 kDa, which matures into a 26 kDa protein after the removal of a 19 AA signal sequence (20, 21). Based on its sequence, OmpV was proposed to share properties with porins (22). Depending on the culture conditions, OmpV can be the most abundant protein in the outer membrane (21, 23–25). Although very little is known about the function of OmpV, a role in pathogenesis has been suggested and antibodies against it can be found in convalescent human sera (21, 26). OmpV is upregulated in *V. parahaemolyticus* in the presence of PmB and in *V. cholerae* in response to human α-defensin 5, while it has a role in osmoregulation in other *Vibrio* species (15, 27). However, its role in resistance to AMPs in *V. cholerae* is yet to be described.

In this study, a transcriptomic analysis of *V. cholerae* grown in the presence of a subinhibitory concentration of PmB showed that the expression of the uncharacterized outer-membrane protein OmpV is increased by PmB. We showed that *vprAB* (*carRS*), *ompV,* and *virK* are organized as a conserved operon in *V. cholerae*, and that its expression is modulated by PmB. The deletion of *ompV* and *virK* led to a higher sensitivity to AMPs, suggesting a role for OmpV and VirK in AMP resistance. The deletion of *ompV* does not lead to membrane destabilization or a reduced sequestration by membrane vesicles. Our results showed that PmB leads to the expression of the RND multi-drug efflux system *vexAB* in an *ompV*-dependant manner. Structural predictions suggest that OmpV is a β-barrel with an unusual membrane-accessible lateral opening that provides entry into a highly electronegative barrel lumen, which could accommodate PmB. This interaction could induce an OmpV conformational change, leading to an intracellular signal event inducing the expression of *vexAB*. We identify OmpV as a new effector for AMP resistance, and the *vprAB-ompV-virK* operon represents the first specialized locus that activates multiple systems for antimicrobial resistance in *V. cholerae*.

## Material & methods

### Strains

*Vibrio cholerae* O1 El Tor strain A1552, an Inaba clinical strain isolated in 1992 from a Peruvian tourist, was used for this study (28). Bacterial strains and plasmids used in this study are listed in Table I. *V. cholerae* strains were grown on LB (10 mg/mL tryptone (Termo Fisher™), 5 mg/mL yeast extract (Thermo Fisher™), 5 mg/mL NaCl) agar plates at 37°C, and cultivated in LB broth at 37°C for 16 h prior to experiments. When needed, L-arabinose (0.2 % w/v) (Thermo Fisher^TM^), carbenicillin (50 µg/mL) (VWR), or PmB (Sigma) were added to the media.

**Table I.** Bacterial strains and plasmids used in this study

| Strain | General characteristics | Reference |
| --- | --- | --- |
| ***V. cholerae*** |  |  |
| A1552 | Wild-type strain, O1 El Tor, pathogenic strain isolated from human cholera infection | (28) |
| A1552Δ*ompV*<br>Δ*virK* | A1552 derived strain in which *ompV* has been replaced by a chloramphenicol resistance cassette, with promoter and terminator, from pKD3, with a knocked down expression of *virK* as determined by qPCR. | This study |
| A1552Δ*virK* | A1552 derived strain in which *virK* has been removed | This study |
| A1552Δ*ompV* | A1552 derived strain in which *ompV* has been removed | Kindly provided by<br>Dr S.N. Wai |
| ***E. coli*** |  |  |
| DH5α | F− φ80*lacZ*Δ*M15* Δ(*lacZYA-argF*)_U169 *recA1 endA1 hsdR17*(r<sub>K</sub><sup>−</sup>, m<sub>K</sub><sup>+</sup>)<br>*phoA supE44 λ-thi-1 gyrA96 relA1* | (34) |
| **Plasmids** |  |  |
| pBAD24 | Expression vector. Arabinose inducible promoter, resistance cassette to carbenicillin (https://www.addgene.org/vector-database/1845/) | (88) |
| pBAD24-*ompV* | pBAD24 vector with the complete annotated *ompV* open reading frame from A1552 under the *ara* promotor | This study |
| pBAD24-*ompV*-153 | pBAD24 vector with the complete annotated *ompV* open reading frame, starting from the ATG in position -153, under the *ara* promotor | This study |
| pBAD24-*virK* | pBAD24 vector with the complete annotated *virK* open reading frame from A1552 under the *ara* promotor | This study |
| pBAD24-*ompV*-153-*virK* | pBAD24 vector with the complete annotated *ompV* open reading frame, starting from the ATG in position -153, annotated *virK* open reading frame and the intergenic region from A1552 under the *ara* promotor | This study |
| pKD3 | Template plasmid for FRT-flanked chloramphenicol resistance cassette (https://www.addgene.org/45604/) | (30) |
| pE-FLP | Vector carrying the flippase FLP used to remove the chloramphenicol cassette flanked by FRT regions (https://www.addgene.org/45978/) | (33) |

### Mutant construction and complementation using pBAD24

The *virK* and *ompV-virK* mutants were obtained as described before using natural competence and PCR products (29). Briefly, the chloramphenicol cassette flanked by FRT regions and P1 and P2 was amplified by PCR from pKD3 using primers adding 50 nt homology (CmR*_ompV_* F/R; CmR*_virK_* F/R) (Table II) to the up- and downstream regions of the target genes (30). Homologous regions of 1000 nt up- and downstream of the target genes were also amplified by PCR using different primers (*ompV*_up_F/R; *ompV*_down_F/R; *virK*_up_F/R; *virK*_down_F/R) (Table II), then linked to the cassette by two-step PCR using *ompV*_up_F and *ompV*_down_R, or *virK*_up_F and *virK*_down_R (31). Two hundred nanograms of the final amplicon were added to A1552 grown for 24 h at 30°C with chitin, in M9 supplemented medium (32). The cells were further incubated for 24 h at 30°C. The mutants were selected on LB agar plates supplemented with 2 μg/ml of chloramphenicol. To remove the resistance cassette, pE-FLP was used as described in (33). The final constructions were verified by PCR and sequencing using *verif* primers (Table II).

**Table II.**
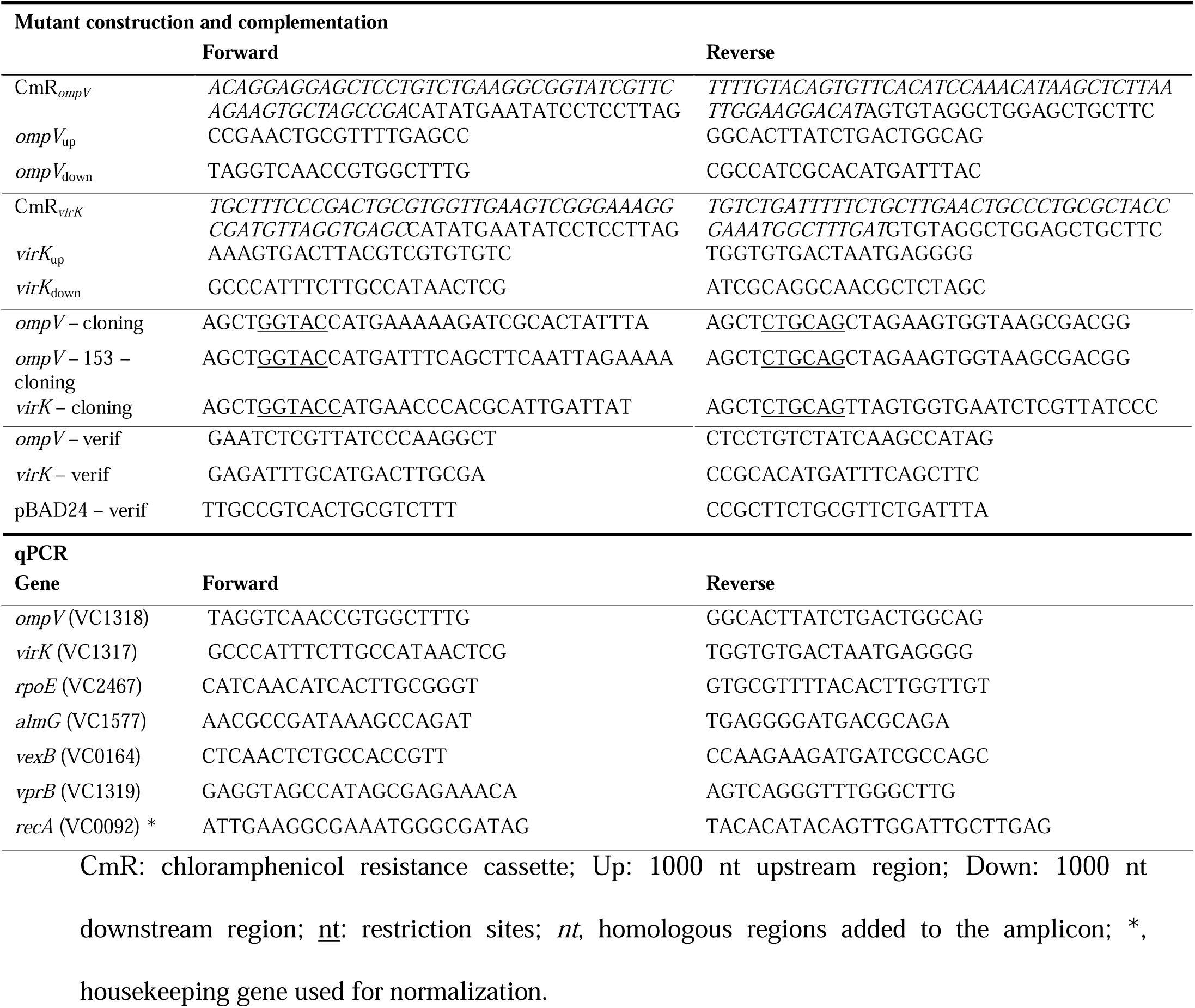
Primers used in this study

The complementation of *ompV* using pBAD24 was carried as described before (29). Briefly, the complete open reading frames (ORF) of *ompV* (<u>VC_1318</u>), from the annotated ATG or the ATG in position –153 pb, and *virK* (<u>VC_1317</u>), were amplified by PCR from A1552 genomic DNA using the primers adding restriction sites listed in Table II. The amplicons and purified pBAD24 vector were digested with PstI and KpnI from New England Biolabs® according to the manufacturer’s instructions, and purified from agarose gel using Monarch Gel purification kit (New England Biolabs®). They were ligated using T4 ligase from New England Biolabs®. The constructions were amplified in thermocompetent *E. coli* DH5α (34), extracted using Pure Yield™ Plasmid Miniprep System (Promega) and electroporated in *V. cholerae* at 1.275 kV, 25 Ω in 1 mm electroporation cuvettes (Thermo Fisher). The vector was maintained with 50 µg/ml carbenicillin.

The *ompV* ORF and its – 153 bp region from other *V. cholerae* strains were compared to those of A1552 using the basic alignment tool nucleotides BLAST® from the National Center for Biotechnology Information (35). The genomes from N16961 (<u>AE003852.1</u>), C6706 (<u>CP046844.1</u>) and MO10 (<u>CP072849.1</u>) were used for comparison. The synteny of the *ompV* region was analysed with PATRIC, using the Compare Region Viewer of the Features section (36, 37) and with SyntTax (38).

### Growth curves and minimal inhibitory concentrations

*V. cholerae* was grown for 16 h at 37 °C with agitation in LB, with carbenicillin when needed. A 1:50 dilution in fresh media was done, and the bacteria were grown at 37°C to an optical density at 600 nm (OD_600nm_) of 0.3. They were further diluted 1:3000 in LB distributed in 96 wells plates with decreasing concentrations of AMP. The bacterial growth was followed by reading the OD_600nm_ every 30 min, at 37°C with agitation. The minimal inhibitory concentration (MIC) was defined as the lowest AMP concentration that inhibits bacterial growth. Data were obtained from at least three independent experiments, in technical triplicates.

### Vesicles extraction, quantification, and visualization on SDS gel

Membrane vesicles (MVs) were isolated from 25 mL cell-free supernatant from 16h cultures in LB, with or without 3 µg/ml of PmB, as described before (39). MVs were suspended in 100 µl of phosphate buffered saline (PBS) and quantified using i) a Bradford assay (Bio-Rad 500-0006) and a bovine serum albumin (BSA) standard curve, as directed by the manufacturer and ii) a fluorescent lipid-labelling FM4-64. Briefly, 2 µL of MVs were incubated with the FM™ 4-64 Dye (N-(3-Triethylammoniumpropyl)-4-(6-(4-(Diethylamino) Phenyl) Hexatrienyl) Pyridinium Dibromide) (Thermofisher) at 2 µg/mL in a final volume of 100 µL in a 96-well black plate. The fluorescence was measured at 515/640 nm using a SpectraMax iD3 reader (Molecular devices). To visualize the MV proteins, 10 µl of the MV preparations were suspended in 20 µl of Laemmli buffer 2X and boiled 10 min at 100°C. Then, 10 µl of the samples migrated on a 13 % sodium dodecyl sulfate gel. Gels were further colored with Coomassie blue.

### Survival in the presence of lethal concentrations of PmB

Ninety microliters of midlog cultures were incubated for 30 min with 500 µg/ml of PmB at 37°C, with or without 5 µl of MV preparations (final concentration 10 X), as described before (39). Ten microliters of ten-fold dilutions were spotted on LB agar plates, then incubated at 37°C for 16 h and numbered. The relative survival was calculated using the number of colony forming units per ml recovered in the presence of PmB in comparison to the non-treated cells. Data were obtained from at least three independent experiments.

### Detection of outer membrane pore formation by fluorescence

Midlog cultures were stained using the fluorescent probes N-phenyl-1-naphthylamine (NPN) and propidium iodide (PI) as described before (40), with modifications. Briefly, NPN and PI were used for the detection of outer membrane and outer/inner membrane damages, respectively. *V. cholerae* was grown to an OD_600nm_ of 1 in LB at 37°C. Bacteria were washed in PBS. Then, PmB at concentration of 50, 25, 10 and 3 µg/ml, or none as control, was added. NPN (20 µM) and PI (20 µM) were added, followed by an incubation of 30 min at room temperature, in the dark. A hundred microliters of each sample were added to a 96-well plate. The fluorescence was acquired with the SpectraMax iD3 reader (Molecular devices) at 350/420 nm and 535/615 nm, respectively. PBS with NPN and PI was used a negative control for autofluorescence to blank the values. The relative fluorescence of each condition was measured in comparison to the wild-type strain without PmB. Data were obtained from at least three independent experiments in technical duplicates.

### RNA extraction, cDNA construction and qPCR analysis

*V. cholerae* was grown to an OD_600nm_ of 0.5 at 37°C in LB, with or without 3 µg/mL of PmB. The bacterial pellets from 10 ml cultures were suspended in 1 mL TRIzol solution (Invitrogen). The total RNA was extracted according to the manufacturer’s instructions and retrotranscribed to cDNA using QuantiTect Reverse Transcription Kit (QIAGEN). Their purity and quality were assessed by nanodrop and migration on 2% agarose gel, respectively. Quantitative PCR analysis was done as described before (39) with primers listed in Table II and using PerfeCTa SYBR® Green FastMix Low ROX (Quantabio). The amplification cycle constitutes of an initial activation step of 30 s at 95°C, followed by 40 cycles of denaturation at 95°C for 5 s, annealing/elongation at 57°C for 17 s and data collection for 12 s at 70°C. The normalized relative expression of various AMP resistance genes was calculated in PmB treated bacteria in comparison to non-treated cells using QuantStudio™ Design and Analysis Software (Thermo Fisher) v1.5.1 and normalized using *recA*. The results were obtained from at least 3 independent experiments, in technical triplicates.

The genomic context of *ompV* was determined using the qPCR primers (Table II) and cDNA of A1552 grown with and without PmB. The intergenic regions between *vprB* and *ompV* and between *ompV* and *virK* were amplified from cDNA by PCR using *vprB*-R and *ompV*-F, and *ompV*-R and *virK*-F. The amplicons migrated on 1 % agarose gel and were visualized with RedSafe™ Nucleic Acid Staining Solution under ultraviolet light.

### RNAseq of total RNA

Total RNA from A1552 grown with or without 3 µg/ml of PmB to an OD_600nm_ of 0.5 was isolated as described in the previous section. The ribosomal RNA depleted RNA was sequenced at the Génome Québec Innovation Center (Centre Hospitalier Universitaire Sainte-Justine, Montréal, QC, Canada). Total RNA was quantified and its integrity was assessed using a LabChip GXII (PerkinElmer) instrument. rRNA was depleted from 125 ng of total RNA using QIAseq FastSelect (−5S/16S/23S Kit 96rxns). cDNA synthesis was achieved with the NEBNext RNA First Strand Synthesis and NEBNext Ultra Directional RNA Second Strand Synthesis Modules (New England BioLabs). The remaining steps of library preparation were done using and the NEBNext Ultra II DNA Library Prep Kit for Illumina (New England BioLabs). Adapters and PCR primers were purchased from New England BioLabs. Libraries were quantified using the KAPA Library Quanitification Kits - Complete kit (Universal) (Kapa Biosystems). Average size fragment was determined using a LabChip GXII (PerkinElmer) instrument. The libraries were normalized and pooled and then denatured in 0.02N NaOH and neutralized using HT1 buffer. The pool was loaded at 175pM on a Illumina NovaSeq S4 lane using Xp protocol as per the manufacturer’s recommendations. The run was performed for 2×100 cycles (paired-end mode). A phiX library was used as a control and mixed with libraries at 1% level. Base calling was performed with RTA v3. Program bcl2fastq2 v2.20 was then used to demultiplex samples and generate fastq reads. Sequencing reads were cleaned with fastp version 0.23.4 and then mapped onto the genomic sequence of *V. cholerae* O1 biovar El Tor strain N16961 (RefSeq <u>GCF_000006745.1</u>) with Bowtie version 2.5.1. After sorting with SAMtools version 1.17, gene-mapping reads were counted with featureCounts version 2.0.1. Finally, differential gene expression was calculated with DESeq2 version 1.40.1 using R version 4.3.0. The output consisted of base mean values, fold change values (Log2(Fold Change)) of genes expression in PmB-treated cells in comparison to non-treated cells, standard error of the estimated fold change values (IfcSE), statistic values (Stat), P values (p-value) and adjusted P values (p-adj) (Table SI). Transcripts with a Log2(Fold Change) < -0.4 or > 0.4, and with a p-adj < 0.05 were considered as significantly modulated by the presence of PmB. The experiment was conducted in biological duplicate. The genes with a modified expression were submitted to the STRING database for network cluster enrichment (41). Sequencing reads from the present project have been deposited in the public SRA database under accession number PRJNA1152934.

### Protein structure prediction and analysis

The signal sequence of OmpV was predicted using SignalP 6.0 (42). The structure of OmpV without its signal sequence was predicted using AlphaFold2 (43) as implemented through ColabFold (44). Protein structure models were visualized using ChimeraX (45). The predicted structure of OmpV was submitted to the Foldseek (46) and DALI (47) servers for comparison to experimentally-determined structures deposited in the Protein Data Bank. The electrostatic surface potential of OmpV was calculated using APBS tools (48) and visualized using ChimeraX. The volume of the putative binding pocket in the lumen of the OmpV barrel was calculated using CASTpFold (49) and visualized using Chimera (50). Structural alignment of OmpV and MipA was performed using the Protein Data Bank pairwise structural alignment tool.

### Statistical Analysis

All data are expressed as mean ± SD and were analyzed for significance using the GraphPad Prism version 10.2.2 for Windows (GraphPad Software, Boston, Massachusetts USA, www.graphpad.com). Student’s *t*-tests were used to compare conditions between 2 groups. Single way ANOVA was used for multiple groups comparison. A result was considered as significant when *p* value < 0.05 (*).

## Results

### A transcriptomic analysis reveals that *ompV* is upregulated in the presence of PmB

To identify new effectors of PmB resistance in *V. cholerae*, a global transcriptomic analysis of the A1552 El Tor strain grown with and without PmB was performed. Cells were grown in the presence or absence of subinhibitory concentrations of PmB. RNA was isolated, then reverse transcribed, and the cDNA was sequenced. Data were analysed for differential expression using the variance analysis package DESeq2. An average number of 70,010,813 reads were identified per sample with an average of 30,597,139 reads being assigned to a gene, and a Q20 > 6.6 representing a base calling accuracy of > 98 % (Table SII). A total of 3715 genes were identified (Figure 1). Genes with a p-adj < 0.05 were considered as significantly modulated by the presence of PmB (colored dots in Figure 1) and are listed in Table SI. Of the 280 modulated genes, 211 and 69 had a significantly increased or decreased abundance of transcripts in the presence of PmB, respectively (Table SI, Figure 1). After the removal of pseudogenes, the identified genes (n=262) with differential expression were submitted for an analysis of network clusters enrichment using STRING (Figure 2) (41). Several clusters were significantly enriched (FDR < 0.05) in the presence of PmB, including 14 genes out of 57 (KEGG pathway: <u>map01503</u>) involved in cationic AMP resistance (Figure 2, Table SIII). The *almEFG* lipid A modification system operon was upregulated in the presence of PmB, as well as the two-component system activating this LPS modification system, *vprAB* (*carRS*), and the RND-transporter *vexAB* (Table SI, Table SIII). The two-component system response regulator VxrB, which contributes to biofilm formation and upregulates the type VI secretion system in response to PmB, also exhibited elevated expression in the presence of PmB (29) (Table SIII). Genes from the type II secretion system cluster, the iron-sulfur binding cluster, multi-drug efflux complex clusters, and the Von Willebrand factor A-like domain superfamily cluster were also significatively enriched (Figure 2). Amongst the genes that were upregulated in the presence of PmB (Table SI), *ompV* stood out because we previously demonstrated that it is also more abundant in the MVs isolated from *V. cholerae* grown in the presence of PmB and the human cathelicidin LL-37 (51). RT-qPCR confirmed that the expression of *ompV* was increased in the presence of PmB by 2.79-fold (Figure 3). Since expression and abundance is elevated in the presence of AMPs, we wondered if *ompV* could be implicated in AMP resistance.

**Figure 1.**
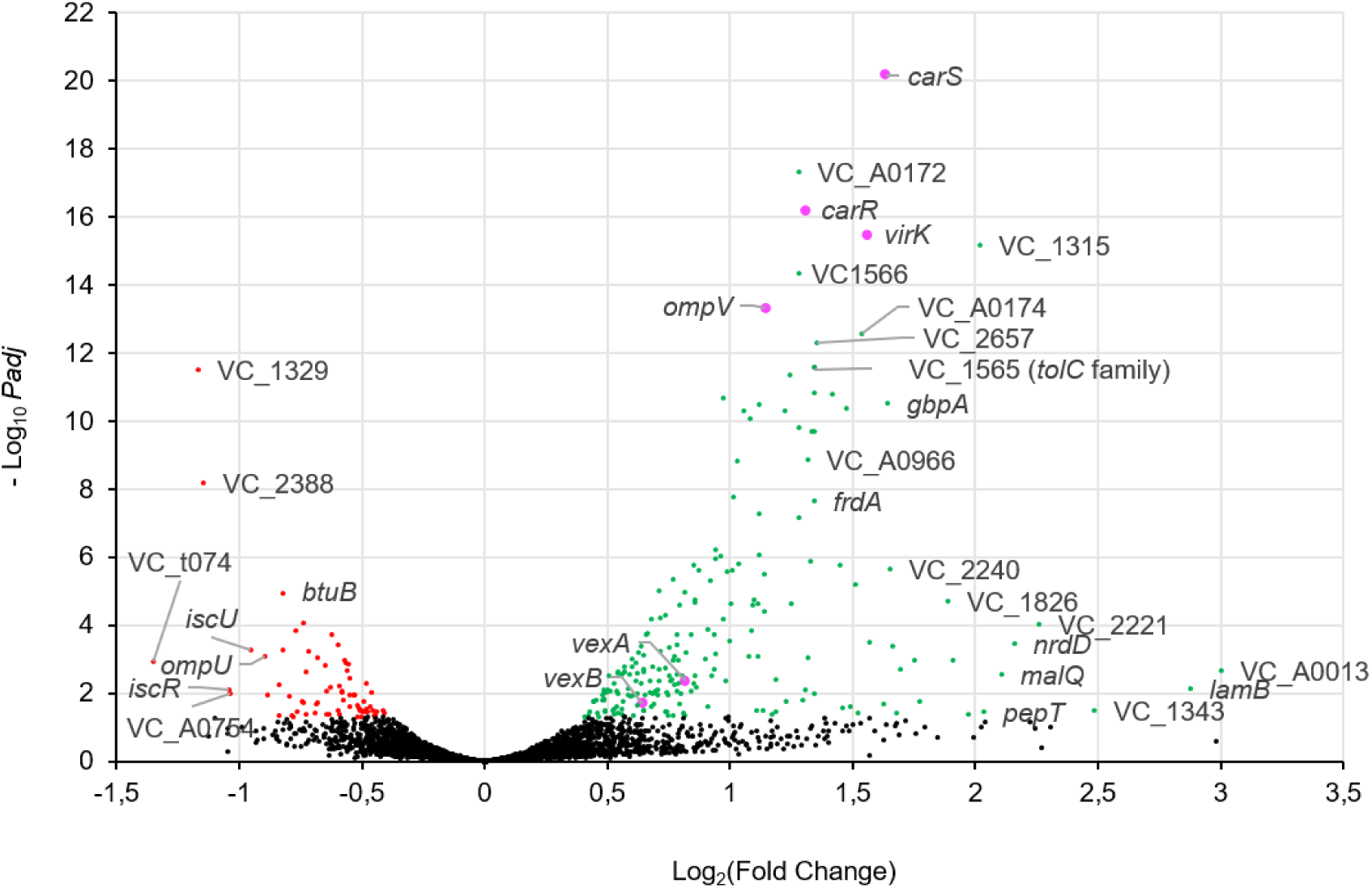
Differential gene expression in *V. cholerae* A1552 in the presence of subinhibitory concentrations of polymyxin B. *V. cholerae* O1 El Tor strain A1552 was grown to midlog phase in LB supplemented with 3 µg/ml of polymyxin B (PmB). The differential gene expression in A1552 treated with PmB in comparison to non-treated cells was analysed using the variance analysis package DESeq2. The expression (Log_2_(Fold Change)) and adjusted P-value (-Log_10_*Padj*) for each identified gene were plotted. A total of 3715 genes were identified. Genes with Padj < 0.05 and Log_2_(Fold Change) < -0.4 or > 0.4 were considered as significantly modulated by the presence of PmB. Black dots represent genes with unmodified expression. Red and green dots represent genes with decreased or increased expression, respectively. Pink dots are genes with increased expression that are the object of this study.

**Figure 2.**
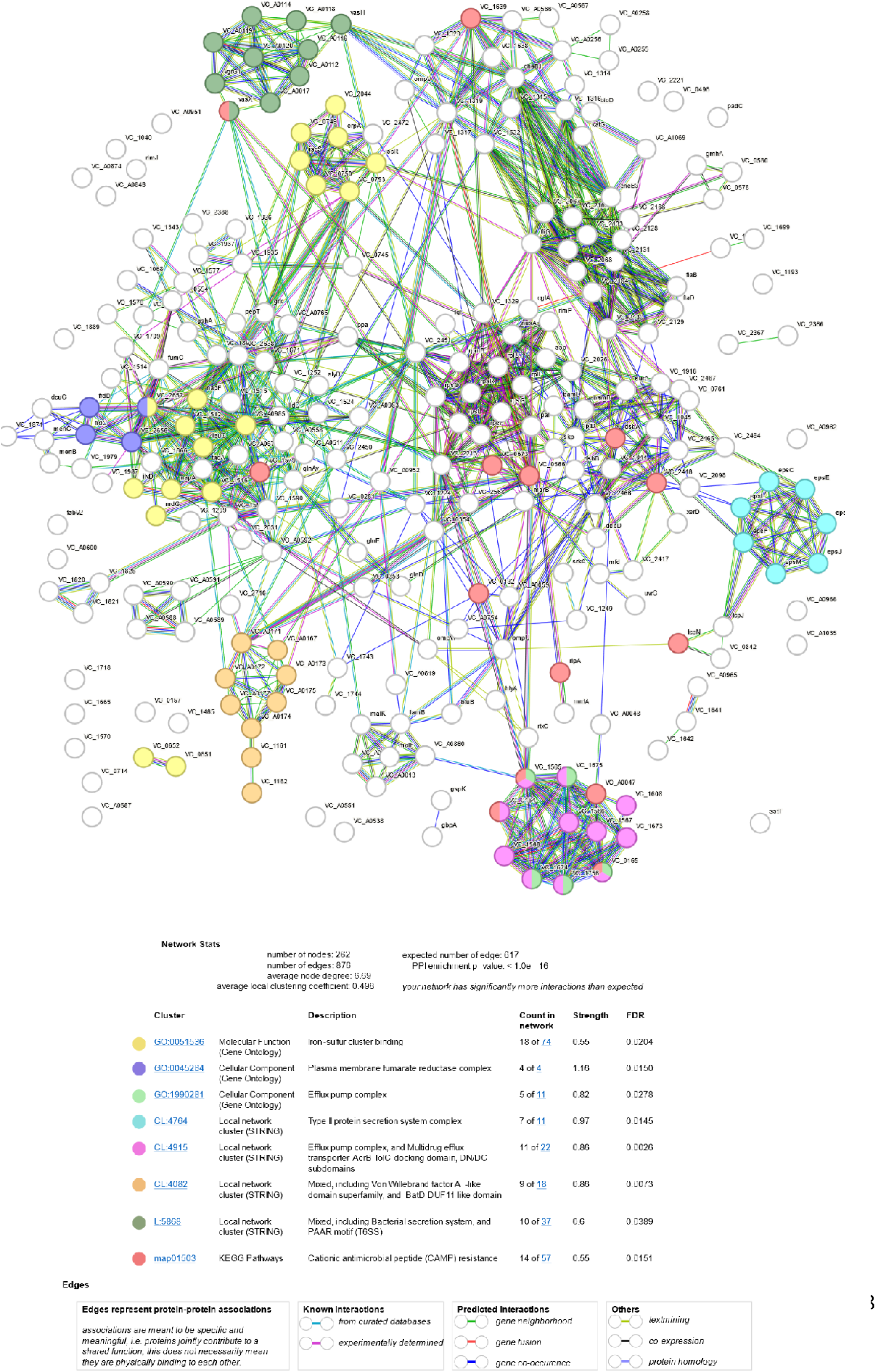
STRING analysis of the genes with modulated expression in the presence of subinhibitory concentrations of polymyxin B. A1552 was grown to midlog phase in LB supplemented with 3 µg/ml of polymyxin B (PmB). The identified genes with significantly modulated expression in the presence of PmB were submitted to STRING for gene cluster enrichment analysis. Colored dots represent significantly enriched (FDR<0.5) clusters of genes with modified expression in the presence of PmB. Edges represent protein-protein associations.

**Figure 3.**
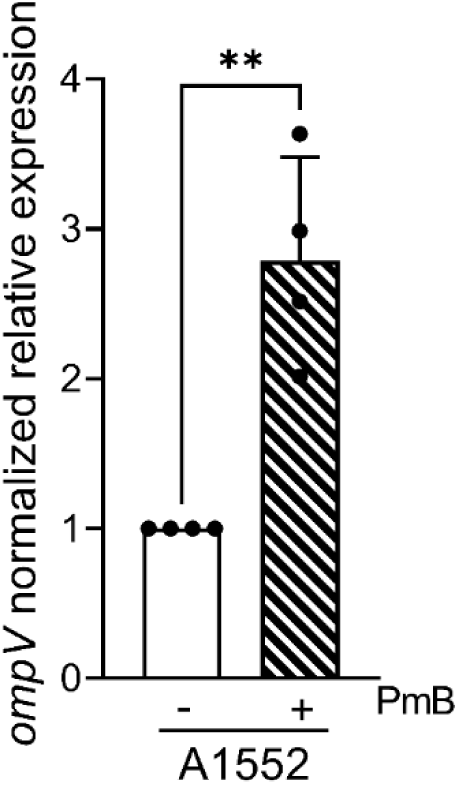
The expression of *ompV* is increased in the presence of polymyxin B in *V. cholerae*. *V. cholerae* A1552 was grown to midlog phase in LB with or without 3 µg/ml of polymyxin B (PmB). The relative normalized expression of *ompV* was determined by quantitative RT-PCR in comparison to untreated A1552 cells, and normalized using *recA*. Data are presented as the mean ± SD from 4 independent experiments conducted in technical triplicates. Asterisks represent a significant difference in expression between treated and non-treated cells, as determined by a single way ANOVA (**, *P* < 0.005).

### The loss of *ompV* increases *V. cholerae*’s susceptibility to antimicrobials

To determine whether the uncharacterized protein OmpV plays a role in resistance to AMPs, an *ompV* isogenic deletion mutant was used (A1552Δ*ompV*). The deletion of *ompV* did not interfere with bacterial growth, as we observed no growth difference between the strains (Figure S1A). The susceptibility of A1552Δ*ompV* and A1552 to AMPs was compared by determination of their minimal inhibitory concentrations (MICs) to PmB and LL-37 (Table III & Figure S1). For A1552, the MICs of PmB and LL-37 were 200 µg/ml. The deletion of *ompV* increased the susceptibility to both AMPs, with a MIC of 100 µg/ml and 12 µg/ml for PmB and LL-37, respectively (Table III & Figure S1BC). Complementation of the *ompV* deletion was first carried out using the pBAD24 vector containing the complete annotated *ompV* (pBAD24-*ompV*). However, the MICs (Table III) were not restored to the wild-type level. Since the presence of OmpV was first observed in the isolated MVs of A1552 grown with PmB (51), the presence of OmpV in the MVs of A1552 pBAD24, A1552Δ*ompV* pBAD24 and A1552Δ*ompV* pBAD24-*ompV* grown with PmB was assessed to confirm complementation (Figure S2). MV crude extracts in denaturing buffer were migrated on SDS-PAGE gels, which were further stained with Coomassie blue (Figure S2). The band corresponding to OmpV is clearly visible in the MVs from A1552 pBAD24 and, as expected, is absent in A1552Δ*ompV* pBAD24. However, it was not restored by the pBAD24-*ompV* construct (Figure S2). To rule out pBAD24-linked expression issues, the native regulatory region of *ompV* (153 nucleotides upstream of the *ompV* start site) in addition to the entire *ompV* ORF was cloned into pBAD24 (pBAD24-*ompV*-153). Using this construct, a band corresponding to OmpV was observed in MV preparations (Figure S2), suggesting that the native regulatory context of *ompV* is important for its expression, as has been observed previously (22). Furthermore, the complementation with pBAD24-*ompV*-153 not only restored the MIC to wild-type levels, but further elevated it for PmB to 250 µg/ml (Table III). Similarly, overexpression of *ompV* in A1552 also resulted in resistance to much higher PmB concentrations, with growth at every tested concentration (Table III). The survival of each strain to a 30 min shock with 500 µg/ml PmB was also assessed and showed that the wild-type strain is nearly 2 times more resistant to this exposure than A1552Δ*ompV* (Figure 5C). Altogether, these results suggest that *ompV* is involved in antimicrobial peptide resistance in *V. cholerae*.

**Table III.** Minimal inhibitory concentrations (MICs) of polymyxin B and LL-37

| Strain | MIC (µg/ml) |  |
| --- | --- | --- |
|  | Polymyxin B | LL-37 |
| A1552 pBAD24 | 200 | 200 |
| A1552 pBAD24-*ompV*-153 | >250 | 200 |
| A1552Δ*ompV* pBAD24 | 100 | 12 |
| A1552Δ*ompV* pBAD24-*ompV* | 100 | 12 |
| A1552Δ*ompV* pBAD24-*ompV*-153 | 250 | 200 |
| A1552Δ*virK* pBAD24 | 200 | 50 |
| A1552Δ*virK* pBAD24-*virK* | 200 | 100 |
| A1552Δ*ompV*Δ*virK* pBAD24 | 100 | 12 |
| A1552Δ*ompV*Δ*virK* pBAD24-*virK* | 100 | 12 |
| A1552Δ*ompV*Δ*virK* pBAD24-*ompV*-153 | 250 | 200 |
| A1552Δ*ompV*Δ*virK* pBAD24-*ompV*-153-*virK* | 250 | 200 |

### The integrity of the membrane is not altered by the loss of OmpV

The integrity of the membrane might be impaired by the loss of *ompV*, one of the most abundant proteins in the outer membrane (22). To determine if the susceptibility of A1552Δ*ompV* to AMPs is due to the destabilisation of the outer membrane, a fluorescent assay was used. *N*-phenyl-1-napthylamine (NPN) is a small molecule that cannot cross the intact outer membrane, but, upon membrane damage, binds to phospholipids and emits fluorescence (52). Propidium iodide (PI) fluoresces once bound to the DNA of bacteria with envelope damage. A1552 and A1552Δ*ompV* were grown to midlog phase, washed in PBS, and permeabilized with increasing concentrations of PmB, up to 50 µg/ml. The bacteria were then labelled with NPN and PI. The relative fluorescence was quantified in comparison to the non-treated wild-type strain in a SpectraMax iD3 plate reader (Figure 4). The fluorescence for NPN increased with PmB concentration, with a maximum fluorescence at 50 µg/ml of PmB for both tested strains (Figure 4A). For PI, the fluorescence was similar at 0, 3, 10 and 25 µg/ml PmB, but was significantly increased for A1552 at 50 µg/ml in comparison to the non-treated bacteria (Figure 4B). This is in agreement with our previous study that showed that PmB treatment does not lead to inner membrane permeabilization, up to 50 µg/ml (53). For both markers, the fluorescence was similar between the strains for a given PmB concentration (Figure 4). These results suggest that PmB produces pores in the membrane similarly for both strains and that the loss of *ompV* has no impact on the integrity of the membrane. Thus, the sensitivity of A1552Δ*ompV* to PmB is not due to permeabilization of the outer membrane.

**Figure 4.**
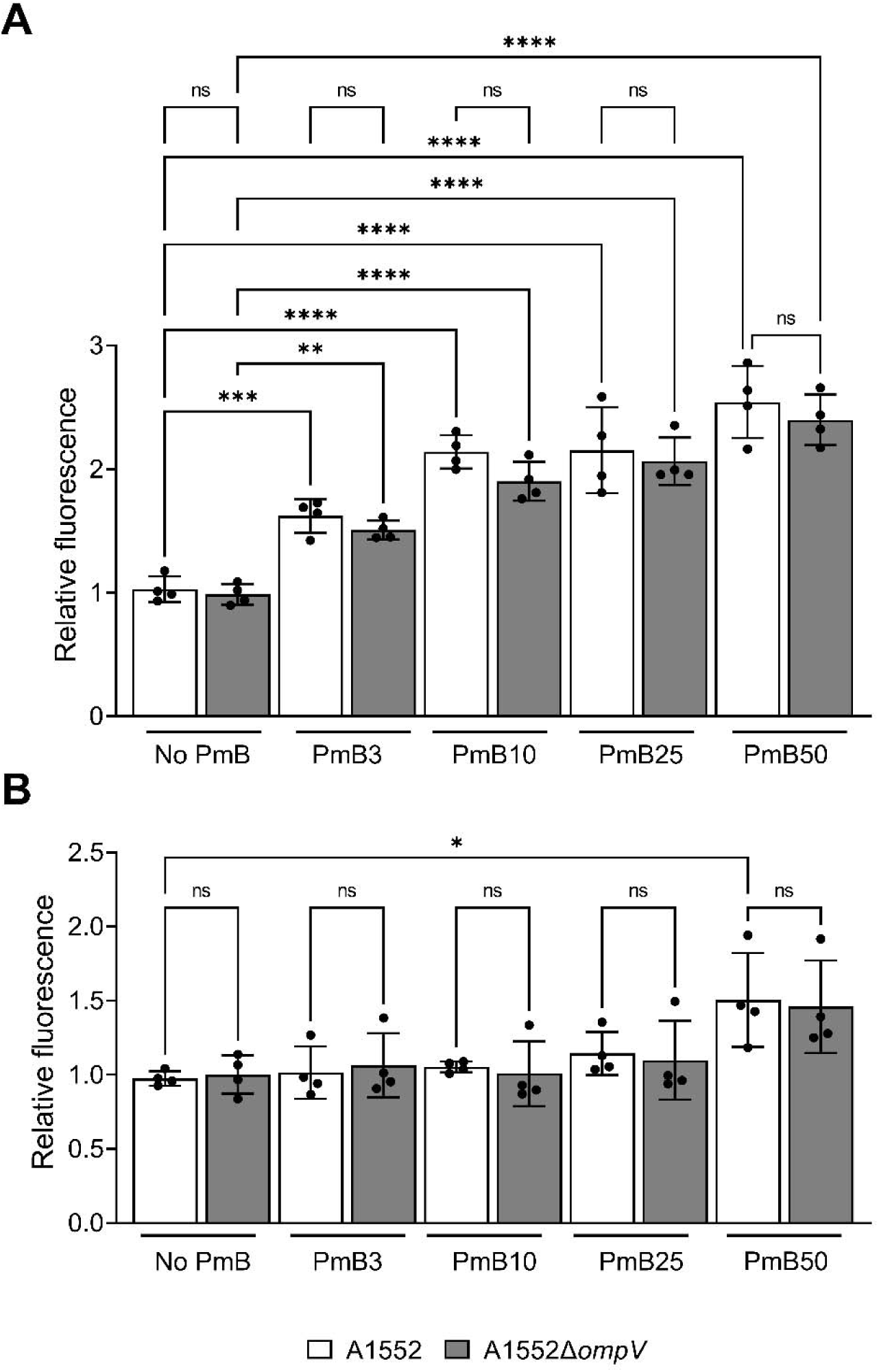
Increased polymyxin B sensitivity of the *ompV* mutant is not due to destabilization of the cellular envelope. Midlog cultures of A1552 and A1552Δ*ompV* were washed in PBS. Then, polymyxin B (PmB) (50, 25, 10 and 3 µg/ml, or none as a control) was added, and the bacteria were incubated for 30 min at 37°C. The bacteria were stained with **A)** *N*-phenyl-1-napthylamine (NPN, 20 µM) and **B)** propidium iodide (PI, 20 µM). A hundred microliters of each sample were added to a 96-well plate and absolute fluorescence was quantified with a SpectraMax iD3 plate reader at 350/420 nm and 535/615 nm for NPN and PI, respectively. The relative fluorescence in each condition was calculated in comparison to A1552 non-treated cells. Data are presented as mean ± SD from 4 independent experiments conducted in triplicates. Asterisks represent a significant difference compared to the non-treated condition within a strain as determined by a single way ANOVA (*, *P* < 0.05; **, *P* < 0.005; ***, *P* < 0.0005; ****, *P* < 0.0001).

### The role of OmpV in antimicrobial resistance is not linked to MVs

When *V. cholerae* is grown with AMPs, the protein content of MVs is modified and greatly enriched in OmpV (51). MVs contribute to antimicrobial resistance by titration and degradation of antimicrobial peptides in the environment (54). We hypothesized that the presence of OmpV in MVs could enhance the titration of antimicrobial peptides, thereby increasing bacterial resistance. If more or bigger MVs were produced in A1552 in comparison to A1552Δ*ompV*, or if the presence of OmpV modified the affinity of the MVs for PmB, then the protection conferred by the MVs of A1552Δ*ompV* would differ from the MVs of A1552. To assess MV production, the isolated MVs from A1552 and A1552Δ*ompV* were quantified using a Bradford assay (Figure 5A) and FM4-64 (Figure 5B), as described previously (39). Both strains produced a similar MV quantity (Figure 5AB), since no significant differences in protein and lipid quantification were observed. This suggests that the sensitivity to PmB upon *ompV* mutation is not due to decreased MV production.

**Figure 5.**
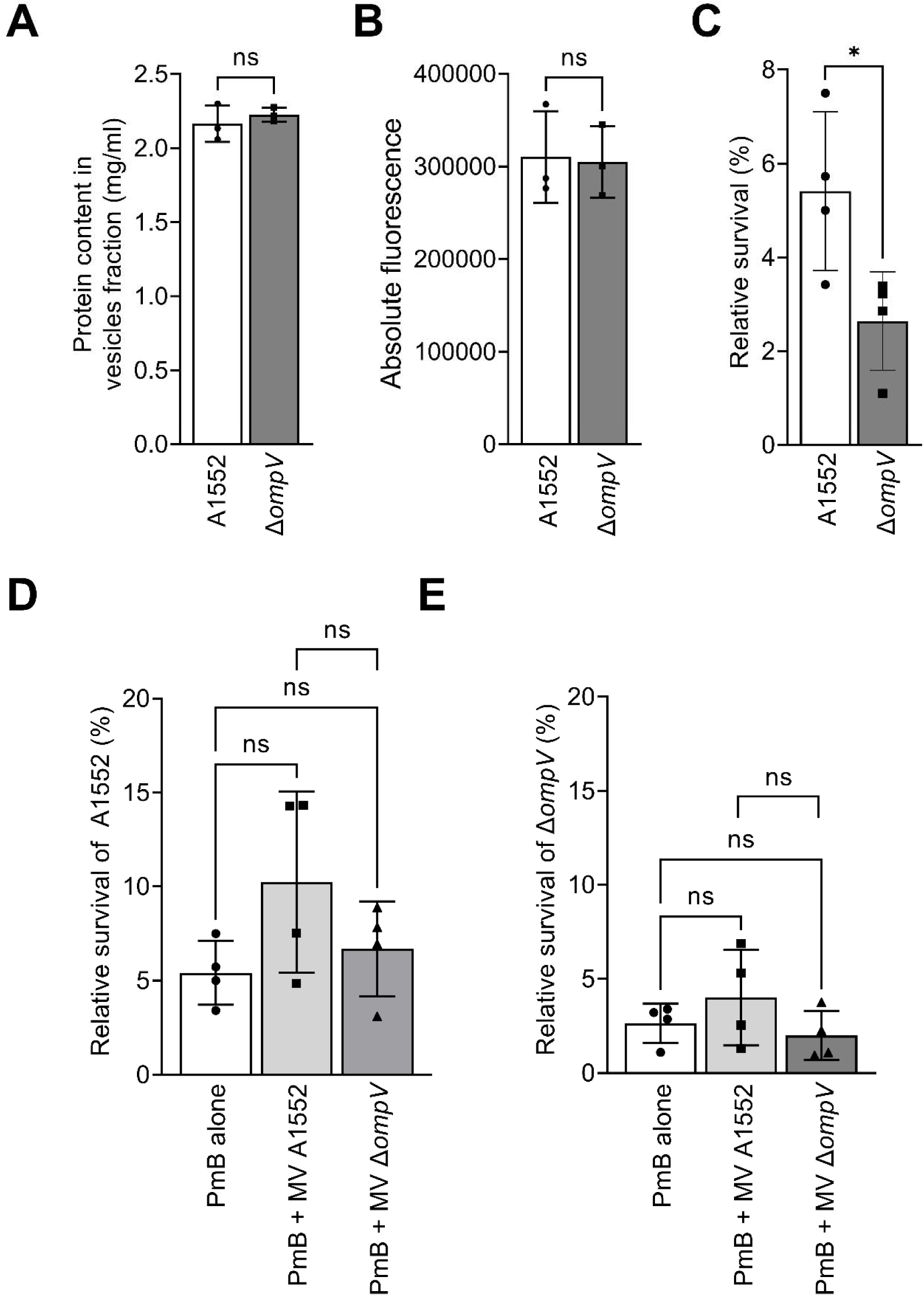
Production of membrane vesicles and their contribution to survival of a 500 µg/ml polymyxin B treatment for A1552 and A1552Δ*ompV*. The membrane vesicles (MV) of A1552 and A1552Δ*ompV* were extracted by ultracentrifugation of the cell-free supernatant from overnight cultures. **A)** The protein content and **B)** lipid fraction were quantified using a Bradford assay and the fluorescent probe FM4-64, respectively. **C)** The survival of A1552 and A1552Δ*ompV* after a 30 min incubation with 500 µg/ml PmB was assessed. The protective effect of the MV from A1552 and A1552Δ*ompV* on **D)** A1552 and **E)** A1552Δ*ompV* to this concentration of PmB was then determined. The surviving bacteria were counted on LB agar. The relative survival was calculated compared to the number of non-treated bacteria retrieved after the incubation. Data are presented as mean ± SD from 4 independent experiments. The asterisk represents a significant difference in survival, as determined by a single way ANOVA (*, *P* < 0.05).

To determine if the affinity of the MVs for antimicrobial peptides is altered upon *ompV* deletion, the effect of the addition of isolated MVs on bacterial survival after a short incubation with 500 µg/ml PmB was then assessed (Figure 5DE). The surviving bacteria were counted on LB agar. As expected, without MVs, the WT strain is more resistant than A1552Δ*ompV*, as more bacteria were retrieved after the incubation (Figure 5C). When MVs from the different strains were added, there was no significant effect on bacterial survival for both strains (Figure 5DE), even though there was a slight increase in survival, suggesting that the presence of OmpV has no or low impact on the capture of PmB by the MVs.

### *ompV* is co-transcribed with the two-component system *vprAB* (*carRS*) and *virK*, and the whole operon is upregulated by PmB

While looking at the genomic context of *ompV*, we noticed that it is located on the minus strand of the first chromosome, clustered with 3 other genes, *vprA* (*carR*)*, vprB* (*carS*), and *virK.* In *V. cholerae*, VirK is an uncharacterized protein, while the two component-system VprAB is responsible for the activation of the LPS modification system *alm* in the presence of PmB (12, 13). This organization is highly conserved amongst *V. cholerae* O1 pre-pandemic, El Tor, and Classical strains, O139 strains, non-O1/non-O139 strains, and some other *Vibrio* species, as determined with PATRIC (https://www.patricbrc.org) (36, 37) (Figure S3). A synteny analysis of the 161 available genomes of *V. cholerae* using <u>SyntTax</u> (38) showed that only 2 strains did not have this arrangement. Although it was found in *V. mimicus* and *V. paracholerae*, this synteny was not found in *V. parahaemolyticus*, *V. vulnificus* and *V. alginolyticus*. To verify if the genes are co-transcribed as an operon on the same mRNA, the intergenic regions between *vprB* and *ompV*, and between *ompV* and *virK*, were amplified by PCR from the cDNA of A1552 using primers inside of the ORF (Figure 6A). Bands corresponding to the expected length were visible on agarose gel, indicating that *vprB*, *ompV,* and *virK* are transcribed on the same mRNA and that they are organized as an operon. The expression of the genes of the *vprAB-ompV-virK* cluster in the presence of PmB was measured by RT-qPCR (Figure 6B). The expression of *ompV* in A1552 grown in the presence of PmB was 2.787 times higher than in the non-treated cells (Figure 6B). The expression of *vprB* and *virK* was 2.515 and 1.986 times higher in the presence of PmB than in the non-treated cells, respectively (Figure 6B). These results suggest that *vprAB*, *ompV,* and *virK* are organized as an operon, and that their transcription is increased by PmB exposure.

**Figure 6.**
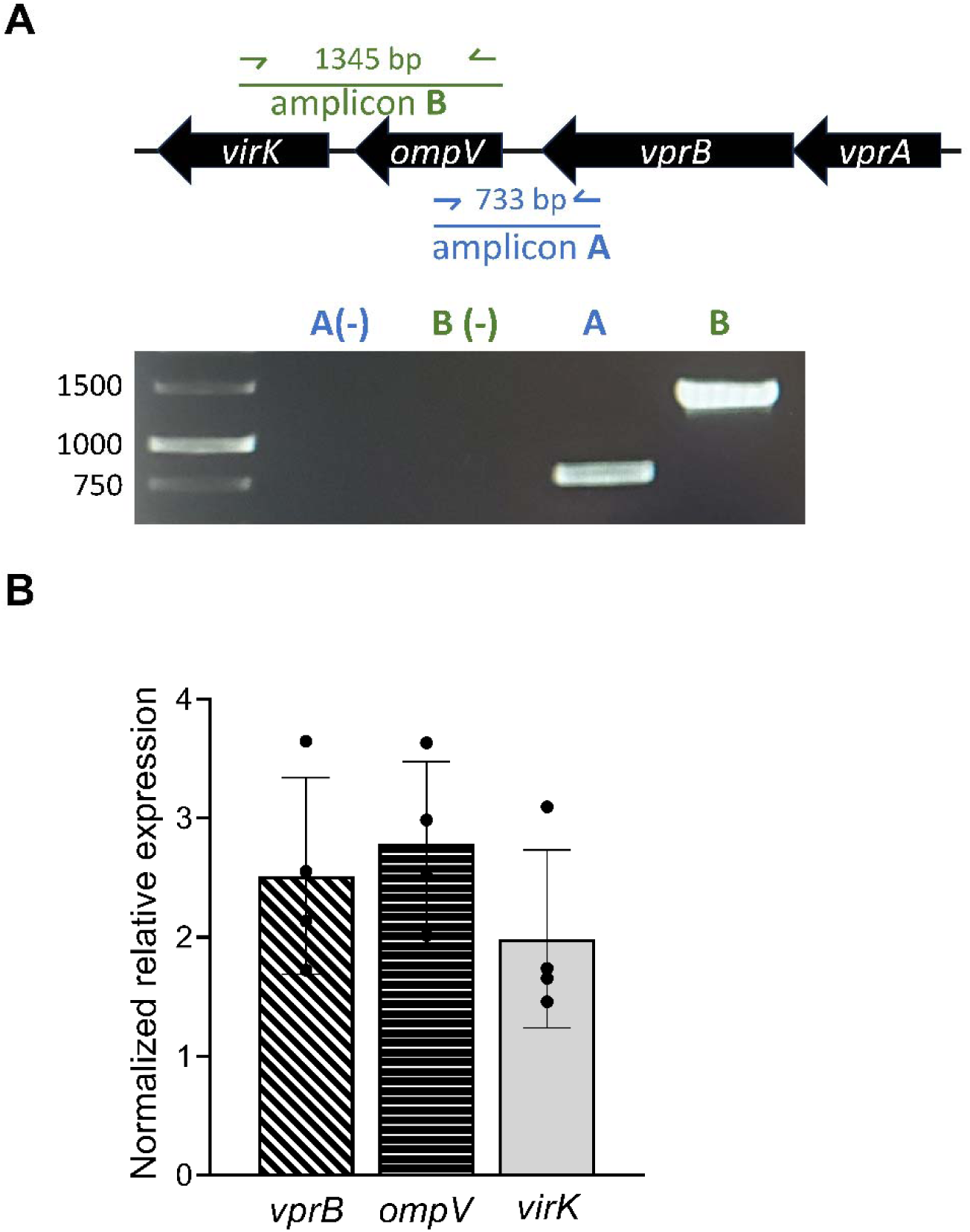
Subinhibitory concentrations of polymyxin B increase the expression of *ompV*, which is organized in an operon with the two-component system *vprAB* and *virK*. A1552 was grown in LB to midlog phase, with and without the addition of a subinhibitory concentration of polymyxin B. Total RNA was extracted and reverse transcribed to cDNA. **A)** Intergenic regions (amplicons A and B) were amplified by PCR using primers (blue and green arrows) in the *vprB*, *ompV,* and *virK* open reading frames, and migrated on a 1 % agarose gel. cDNA was replaced with water in PCR reactions as a negative control (-). The results are representative of 3 independent experiments. **B)** The normalized relative expression of *vprB*, *ompV,* and *virK* was measured in polymyxin B treated bacteria in comparison to non-treated cells by quantitative PCR. The expression was normalized using *recA*. The results are presented as mean ± SD and were obtained from 4 independent experiments, in technical triplicates.

The role of *vprAB* (*carRS*) in resistance is already known (13, 55) and we have demonstrated here that *ompV* plays a role in antimicrobial resistance. To determine if *virK* is also implicated in AMP resistance, a knock-out mutant (A1552Δ*virK*) and a complemented strain (A1552Δ*virK* pBAD24-*virK*) were constructed. Their MIC to AMPs was determined, which showed that A1552Δ*virK* was more sensitive to LL-37, and that complementation partially restored the phenotype (Table III). Although the MICs for PmB were similar for the wild-type strain and A1552Δ*virK* (Table III) the growth of A1552Δ*virK* with 100 μg/mL of PmB clearly showed a defect in comparison to A1552 (Figure S1). These results indicate that A1552Δ*virK* is also more sensitive to AMPs. To determine if there is an additive effect of *ompV* and *virK* deletion on antimicrobial resistance, a A1552Δ*ompV*Δ*virK* double mutant and a complemented strain (A1552Δ*ompV*Δ*virK* pBAD24*-ompV-virK*) were constructed. The MICs of PmB and LL-37 of the double mutant were similar to that of A1552Δ*ompV* (Table III), which were lower than those of A1552Δ*virK*. A single *ompV* or a double *ompV-virK* complementation, but not a single *virK* complementation, led to a total restoration of the MIC in A1552Δ*ompV*Δ*virK* (Table III). These results suggest that there is no additive effect of the double mutation on the sensitivity of the strains to AMP.

### The deletion of *ompV* modified expression of the antimicrobial resistance related gene *vexB*

Previous studies have demonstrated that outer membrane proteins, such as OmpU in *V. cholerae*, can sense and signal for the presence of AMPs (56). This ability allows the bacteria to trigger resistance mechanisms by modifying the LPS and its electrostatic affinity for cationic molecules (57). OmpV may play a similar sensing role for AMPs. In the transcriptomic analysis, we observed that several known resistance effectors of *V. cholerae* were upregulated in the presence of PmB (Table SI and SIII). They include the two-component system *vprAB* (*carRS*) responsible for activating transcription of the *alm* operon (13), the glycyltransferase *almG* that modifies lipid A (6), the RND efflux pump *vexAB* (16), and the alternative sigma factor *rpoE* (56) (Table SI). To determine if OmpV is involved in the regulation of these effectors, a quantitative RT-PCR analysis was conducted in the absence and presence of PmB in both A1552 and A1552Δ*ompV* (Figure 7). As expected, a strong and significant upregulation of *almG*, *vprB*, *rpoE,* and *vexB* in the presence of PmB in A1552 was observed (Figure 7). While *vprB* expression was increased in the presence of PmB in the A1552Δ*ompV* strain, its upregulation was lower than in A1552 (Figure 7A). This suggests that *vprAB* over-expression in the presence of PmB is partly dependent on *ompV*, although it had no effect on the expression of *almG*, part of the VprA regulon (Figure 7B). The expression of *vexB* was not increased by the presence of PmB in A1552Δ*ompV* (Figure 7D). This suggests that the upregulation of the efflux-pump *vexAB* depends on the presence of *ompV*, and that the increased sensitivity of the A1552Δ*ompV* strain could be due to the lack of *vexAB* upregulation. Taken together, these results suggest a role for OmpV in the transcriptional regulation of PmB resistance genes.

**Figure 7.**
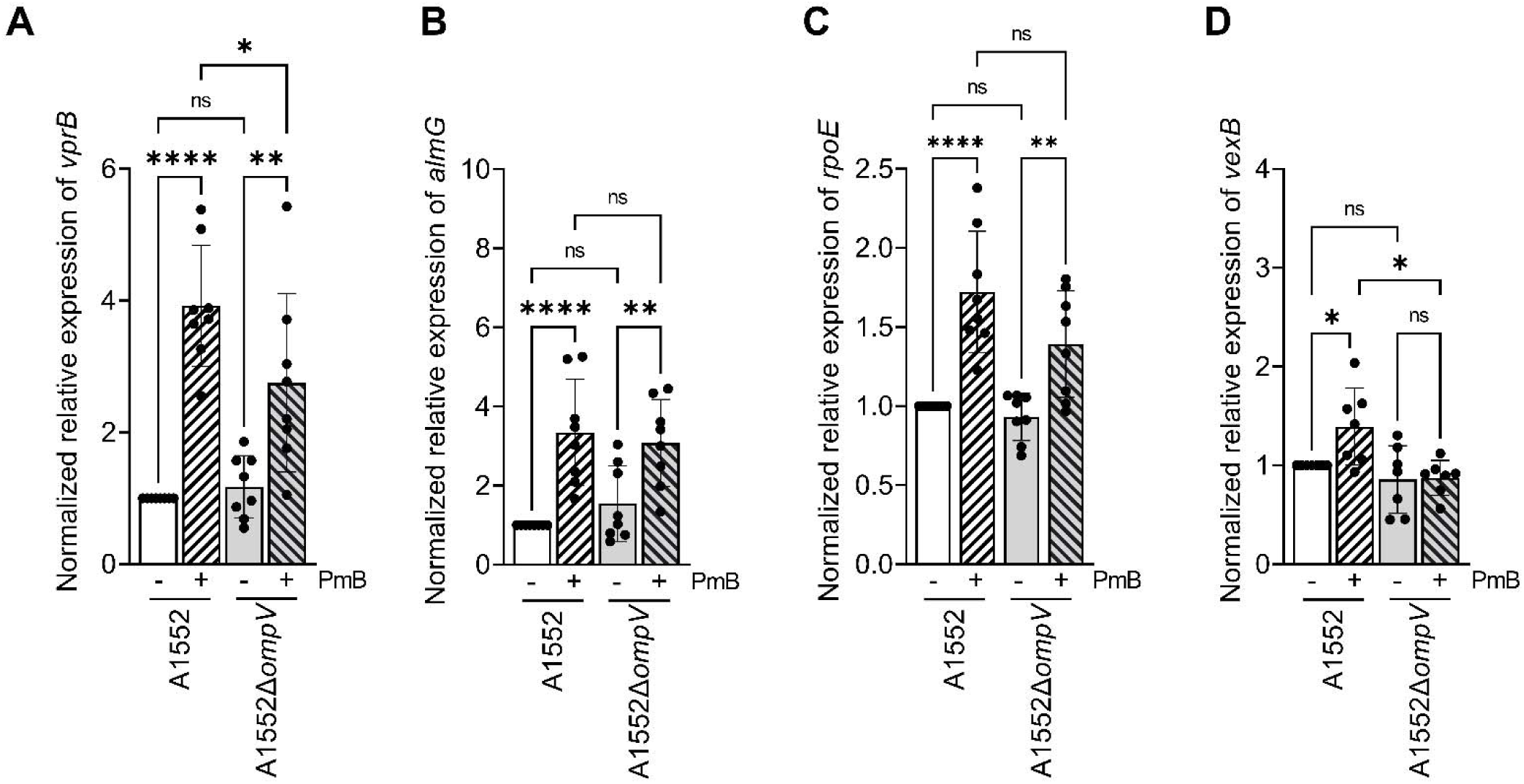
*ompV* deletion modifies the expression of antimicrobial resistance related genes in the presence of polymyxin B. A1552 and A1552Δ*ompV* were grown to midlog phase in LB with or without 3 µg/ml of polymyxin B (PmB). The relative normalized expression of **A)** *vprB,* **B)** *almG*, **C)** *rpoE,* and **D)** *vexB* was determined by quantitative RT-PCR in comparison to the A1552 non-treated cells and normalized using *recA*. Data are presented as mean ± SD from 8 independent experiments conducted in technical triplicates. Asterisk represents a significant difference in expression between treated and non-treated cells, as determined by a single way ANOVA (*, *P* < 0.05; **, *P* < 0.005; ***, *P* < 0.0005; ****, *P* < 0.0001).

### OmpV has a membrane-accessible lateral opening into an electronegative pocket in the β-barrel lumen

To gain further insight into how OmpV may function in the process of AMP resistance, we predicted the structure of OmpV, without its predicted signal sequence (residues 1-19) using Alphafold2 as implemented within ColabFold (44). This suggested that OmpV adopts a 12-stranded β-barrel fold (Figure 8A and S4A). However, β-strands five and six, which form part of the lateral wall of the barrel, do not fully span the membrane-embedded region of the protein (Figure 8A). These short β-strands leave a gap in the barrel wall that provides lateral access to the β-barrel lumen of OmpV (Figure 8A and B). Submission of the predicted structure of OmpV to the Foldseek (46) and DALI (47) servers suggested that there are no experimentally determined structures of β-barrels in the Protein Data Bank with a similar lateral opening. Given the positioning of this gap in the membrane-spanning region of OmpV (Figure 8C), it is likely that it is only accessible through the lipid bilayer of the OM. The opening in the lateral wall of the OmpV barrel leads to a pocket in the lumen that is isolated from both the extracellular and periplasmic spaces due to the presence of several extracellular loops and a periplasmic plug region, respectively (Figure 8A and C). This pocket is highly electronegative (Figure 8B) and has a volume of 1623 ^3^ as determined by CASTpFold (49) (Figure 8C), therefore it could accommodate cationic AMPs such as PmB, which has an approximate volume of 1047 ^3^ based on its structure in complex with human lysine-specific demethylase 1 (PDB 5L3F). Such an interaction could induce conformational changes in OmpV that serve as a signal to initiate cellular processes that protect against cationic AMPs.

**Figure 8.**
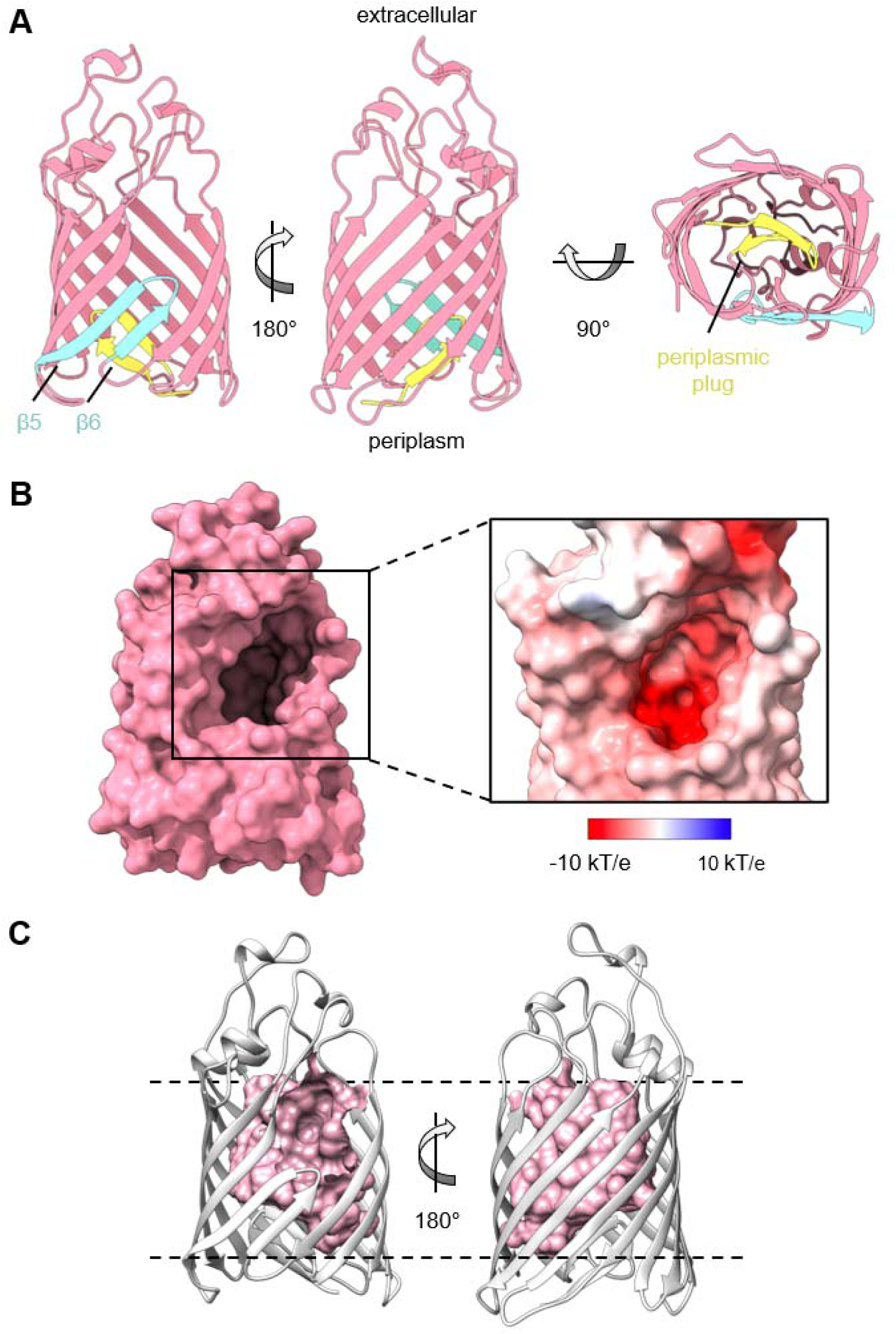
OmpV has a membrane-accessible lateral opening into an electronegative pocket in the β-barrel lumen. **A)** The structure of OmpV without its signal sequence (residues 1-19), predicted by Alphafold2 as implemented within ColabFold. The side of OmpV that faces the periplasmic or extracellular space is indicated. β-strands five and six (β5 and β6), which are shorter than the other strands that comprise the barrel, are indicated in blue. The N-terminal plug region, which occludes access to the barrel lumen from the periplasmic side, is indicated in yellow. **B)** (left) Space-filling model of the predicted structure of OmpV, highlighting the lateral opening in the side of the barrel. The model is oriented similar to the leftmost structure of panel A. (right) Magnified view of the lateral opening, looking directly into the pocket in the barrel lumen. The surface is colored according to electrostatic potential calculated using APBS; contoured from -10 kT/e (red) to 10 kT/e (blue). **C)** The pocket in the lumen of the OmpV barrel as determined by CASTpFold, depicted as a negative space-filling surface in pink. The dashed lines indicate the approximate location of the membrane-solvent boundary.

## Discussion

In this study, we identified the outer-membrane protein OmpV as a new effector of antimicrobial resistance in *V. cholerae* by coupling transcriptomic analyses of knockout mutants with AMP susceptibility assays. We showed that *ompV* is part of a specialized 4-gene operon that activates multiple resistance factors in response to PmB, *i.e*. the LPS modification system *almEFG* and the efflux pump *vexAB*. A structural analysis also determined that OmpV could act as a sensor for PmB in the outer membrane. This operon is widespread in *V. cholerae*, suggesting that this may represent a generalizable strategy for AMP resistance in this species.

Even at a concentration well below the MIC, PmB significantly impacted the transcriptome of *V. cholerae*, with more than 280 genes whose expression was significantly modulated by the presence of PmB. The PmB concentration used (3 μg/ml) is below that required for pore formation and is 1.5 % of the MIC (53). In our previous proteomic analyses, a similar number of proteins (n=241) with a modified abundance in the presence of this PmB concentration were identified (29, 51, 58). This included many genes involved in antimicrobial resistance, such as the main locus responsible for resistance in El Tor strains, *almEFG,* in addition to *vxrB, vprB* (*carS*)*, vexAB,* and the σ^E^ regulatory protein RseB (29, 58). Both analyses also identified that the type II and VI secretion systems were upregulated in the presence of PmB, as well as components of the flagellum. Many RND-transporters and efflux systems were upregulated in the presence of PmB, as would be expected since RND-transporters are responsible for basal resistance to antibiotics and antimicrobials, and because they are regulated by the presence of their substrates (59–61). Other transcriptomic analyses of *V. cholerae* El Tor strains C6706 and El2382 grown in the presence of AMPs have been performed (15, 62). In those analyses, components from the *alm* operon were also upregulated, as well as *vprAB* (*carRS*), transporters, and efflux systems. Interestingly, the studies in strains C6706 and El2382 showed that *ompV* was upregulated in the presence of AMPs (15, 62), but without attributing a role for it in antimicrobial resistance. The fact that *ompV* is highly expressed and abundant in *V. cholerae* in the presence of multiple AMPs from different families caught our attention and made us wonder about its role in antimicrobial resistance.

OmpV is a major protein of the outer membrane of *V. cholerae* (22). OmpV plays a role in adhesion and invasion of enteric epithelial cells in *Salmonella* (63, 64), and is an osmotic stress responsive protein in the marine bacteria *Photobacterium damselae*, *V. alginolyticus,* and *V. parahaemolyticus* (65–67). In those *Vibrio* species, *ompV* has a different genomic context, and is not clustered with *vprAB* and *virK*. OmpV is annotated as a protein of the MltA-interacting protein (MipA) family (68), which is implicated in antimicrobial resistance in *E. coli* (69). Although a role for OmpV in pathogenesis has been suggested in *V. cholerae* (21, 22, 26), its function has not been identified so far in this bacterium. We previously showed that OmpV is found in abundance in MVs of *V. cholerae* A1552 grown with AMPs (51). Here, a transcriptomic analysis and quantitative RT-PCR confirmed that *ompV* is indeed upregulated in the presence of PmB. MIC values and relative survival to a short incubation time with a high PmB concentration showed an increased sensitivity to AMPs upon *ompV* mutation, demonstrating a role for OmpV in resistance to antimicrobials.

The genomic context of *ompV* in *V. cholerae* showed that it is part of a four-genes cluster (*vprA-vprB-ompV-virK*) on the first chromosome, and that the synteny of this cluster is conserved amongst some *Vibrio*. We demonstrated that these genes were transcribed as an operon, and that the whole operon is upregulated by the presence of PmB. It has previously been demonstrated that the expression of *ompV* and *vprB* increased in the presence of human α-defensin 5 (15) and PmB (62). VprAB (CarRS) is a two-component system in which VprB is a sensor histidine kinase, sensing AMPs in the periplasm, and VprA is the response regulator that activates the *almEFG* mediated LPS modification system in the presence of PmB to decrease its interaction with the membrane (12, 13, 15). In this study, we showed that OmpV and VirK are also implicated in antimicrobial resistance.

Our RT-qPCR analysis of known PmB-resistance factors demonstrated that *ompV* is important for the upregulation of the RND-transporter *vexAB* in the presence of PmB. There are six RND-transporter systems encoded in the *V. cholerae* genome that work synergically to provide resistance to various toxic substrates such as bile, detergents, and antimicrobials (16, 19, 70, 71). All of them form a complex with the outer-membrane pore protein TolC to span both the inner and outer-membrane (19, 72). A knock-out mutant of all the RND systems led to a reduction in cholera toxin and toxin-coregulated pilus production, and was attenuated in an infant-mouse model of infection (19, 73). The VexAB RND-system (VC_0164, VC_0165), in which VexA is the membrane fusion component and VexB is the RND pump-protein, is regulated by bile acids and is induced during mammalian intestinal colonisation (16, 74). This system is very similar to AcrAB-TolC of *E. coli* (75). Together with VexCD, VexAB contributes to bile resistance, while the deletion of *vexB* alone leads to a higher susceptibility to SDS, Triton X-100, erythromycin, novobiocin, and PmB, but not to other antibiotics such as β-lactams, aminoglycosides, fluoroquinolones, and tetracyclines, amongst others (16, 19). VexAB requires the *tetR*-family transcriptional regulator *vexR*, which is located upstream of *vexAB* in the same operon, for activation upon substrate exposure (60). *vexR* was upregulated 1.28-times in the presence of PmB in our transcriptomic analysis, although this was not significant (p-adj = 0.1145).

There are five TolC homologues in *V. cholerae*’s genome (VC_1409, VC_1565, VC_1606, VC_1621, VC_2436), with VC_2436 exhibiting the highest similarity to *E. coli* TolC (72). *In vitro*, only the deletion of VC_2436 affected antimicrobial resistance, including to PmB, which was similar to a mutant in which the six RND-systems were deleted. Therefore, VC_2436 is likely to be the outer-membrane pore used by RND-transporters in *V. cholerae* (19, 72, 76). In our transcriptomic analysis, *tolC* (VC_2436) was not upregulated by the presence of PmB. However, because the inner membrane components of RND-systems are generally regulated by their efflux substrate concentrations, the expression of *vexB*, but not of *tolC*, was increased by the presence of PmB, which could explain why the expression of *tolC* was not modified by PmB in our analysis (59, 76). Although VC_1565 was upregulated by PmB in C6706 and in our analysis, VC_2436 was not upregulated eighter in the presence of AMPs in EL2382 and C6706 in the transcriptomic analyses (15, 62), suggesting that *tolC* might be regulated differently than the inner components of the RND systems.

Since the modulation of *vexAB* upon PmB stimulation is *ompV*-dependent and OmpV is an outer membrane protein, we wondered how this response could occur. Some outer-membrane proteins can sense extracellular signals and induce bacterial adaptation to stresses, such as OmpU (56, 57, 77). OmpU signaling is dependent on the action of the envelope-stress response mediated by the alternative σ^E^factor (RpoE) and the proteases DegS, RseA, and RseP (57, 78). In the absence of stress conditions, σ^E^ is segregated at the inner membrane by the anti-sigma factor RseA, preventing its activity on RNA-polymerase (79). The accumulation of misfolded OmpU containing the C-terminal YxF motif leads to the activation of DegS (78, 80). The activated DegS cleaves RseA, which is further cleaved by RseP located in the inner membrane, leading to the release of σ^E^ and to the transcription of the envelope-stress response genes (81, 82). In the presence of AMPs, the disruption of the outer membrane could block the insertion of OmpU in the membrane, leading to the accumulation of mislocalized OmpU in the periplasm, or the binding of AMPs to OmpU could induce a conformational change, promoting exposure of the YxF motif, thus leading to the release of σ^E^ and transcription of membrane-repair genes (57). OmpV has the OMP-periplasmic-stress associated YxF motif. It is thus possible that OmpV also acts as a sensor of AMPs and signals their presence at the outer membrane to activate a transcriptomic regulator inside the cell and the expression of resistance genes including *vexAB*. A study using deletion mutants of different OMPs showed that the outer-membrane activation of σ^E^ by YxF is OmpU-dependant (83). Our qPCR results show that σ^E^ is sill activated upon PmB stimulation in a A1552Δ*ompV* mutant, suggesting that another signal is used. A possible explanation for the modulation of *vexAB* in the presence of PmB is that it could be part of the *vprA* regulon, especially because *ompV* and *vprAB* belong to the same operon. Our quantitative RT-PCR analysis showed that the expression of *vprB* was significantly upregulated in A1552Δ*ompV* in the presence of PmB, but that this increase was also significantly lower than in A1552. In *V. vulnificus*, upon PmB stimulation, *vprA* (*carR*) activates the expression of *eptA* and *tolCV2*, an LPS modification system and a tripartite efflux-pump, respectively (55, 84). However, the expression of *almG*, which is a known effector of the *vprA* regulon (12), is not reduced in A1552Δ*ompV*. The transcription of *vprAB* (c*arRS*) in the presence of PmB could be activated by multiple pathways, and thus only partly depend on *ompV*. It could also mean that lower expression levels of *vprAB* are sufficient to strongly activate the transcription of the *alm* operon, but not *vexAB*, as the affinity of VprA to their respective promotors could be different.

We wondered how OmpV could act as a sensor for PmB and so examined its predicted structure. This revealed that it adopts a β-barrel architecture where both its periplasmic and extracellular openings are occluded. Instead, two short β-strands in the lateral wall of the OmpV barrel create an opening into an electronegative pocket in the lumen of the barrel that is accessible only through the membrane. The proposed porin function of OmpV is questioned (66, 85). We propose that, instead of functioning as a porin, OmpV may directly bind PmB via this electronegative pocket, thus serving as a sensor.

According to the structure prediction and electrostatic surface potential calculations, we propose a model in which OmpV, a MipA structural ortholog (Figure S4B), can sense PmB when it integrates into the outer-membrane via direct interaction with the electronegative barrel lumen, accessible through a membrane-integral gap in the β-barrel wall, and activate the expression of the efflux-pump VexAB through an unknown mechanism (Figure 9). While VirK is predicted to be a cytoplasmic protein, an *E. coli* ortholog similarly predicted to localize to the cytoplasm has been shown experimentally to localize to the periplasm (86). It is therefore possible that VirK could be a periplasmic mediator involved in signal propagation of PmB-sensing by OmpV, especially since they are co-transcribed as part of the same operon. An OmpV conformational change upon PmB binding could thus signal intracellularly using VirK as an intermediary. This signalling might occur through the VprAB system to activate the expression of the *vexAB* efflux pump (Figure 9). VprB is known to respond to cationic AMPs, but the direct interaction between PmB and VprB is yet to be confirmed. During the preparation of this manuscript, a similar system was identified in *Pseudomonas aeruginosa* in which the OmpV ortholog MipA functions as a PmB-sensor, inducing a MipA conformational change that releases the periplasmic mediator MipB (87). This leads to the expression of the efflux pump MexXY through the ParRS two-component system (87), suggesting a conserved mechanism of AMP signaling and resistance. However, in *P. aeruginosa*, the system is not encoded as a single operon and is present in the bacterial genome only in the absence of the *arn* operon, responsible for LPS modification (87). In *V. cholerae*, we have identified a single conserved operon that regulates multiple antimicrobial resistance mechanisms, specifically the efflux pump VexAB and the LPS modification system *alm*. To our knowledge, no other operon encoding multiple resistance system regulators has been identified so far. Future studies are necessary to dissect the role of VirK in antimicrobial resistance, as well as to confirm the involvement of VprAB in *vexAB* regulation or to identify the regulators of the OmpV-mediated response.

**Figure 9.**
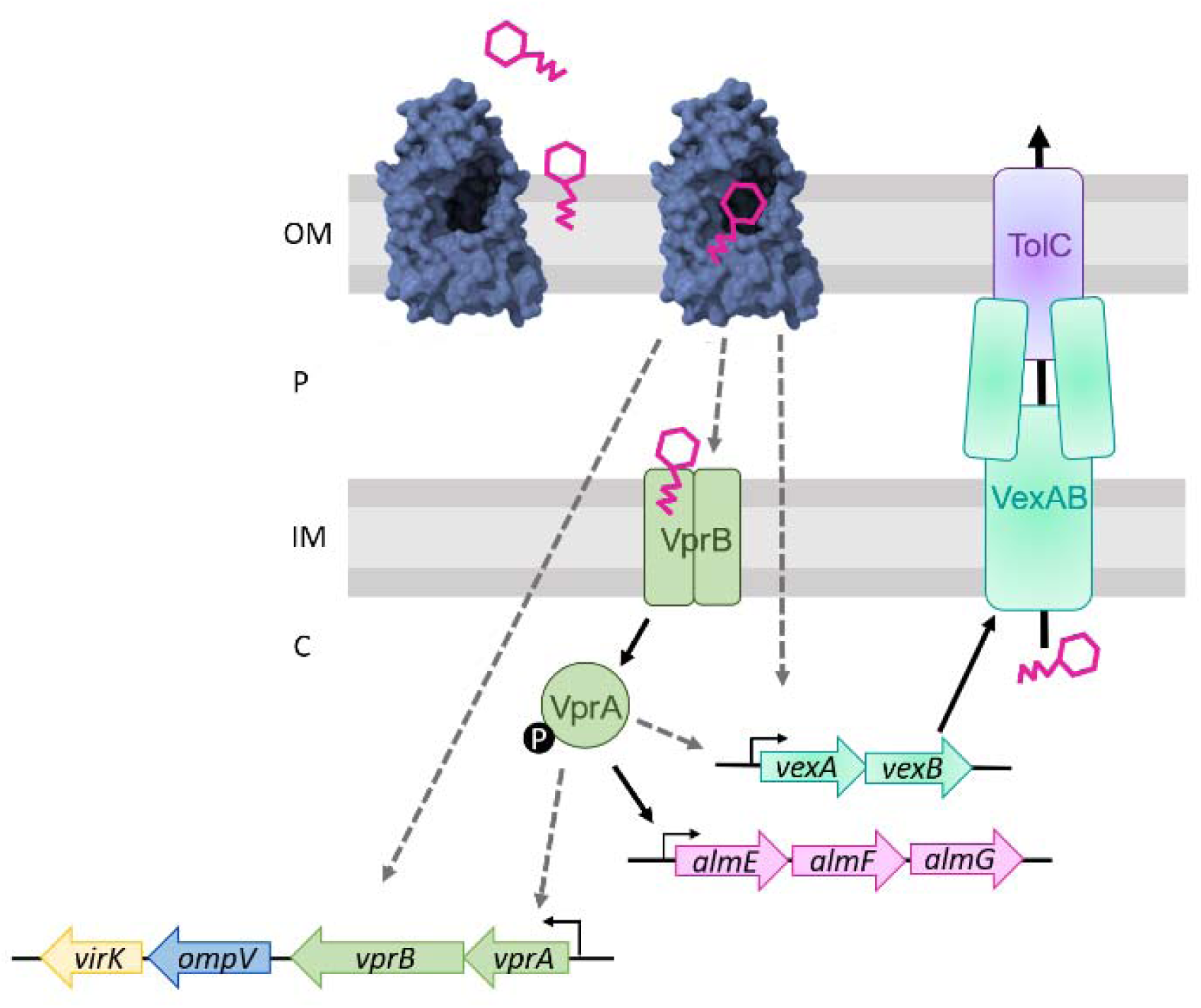
Graphical summary of the proposed mechanism of resistance to polymyxin B mediated by the *vprA-vprB-ompV-virK* operon. OmpV (blue) at the outer-membrane (OM) serves as sensor for polymyxin B (pink). Once polymyxin B integrates into the OM, it can access the electronegative lumen of OmpV through a lateral gap in the barrel wall. This interaction could induce a conformational change, leading to an intracellular signaling event. Using unknown mediators in the periplasm (P) and cytosol (C), or through the two-component system VprAB, the expression of the *almEFG, vprAB-ompV-virK,* and *vexAB* loci is induced. VexAB, together with TolC, forms an efflux-system implicated in resistance to polymyxin B. Known activation pathways are identified using solid black arrows. Hypothesized pathways are represented with dashed grey arrows. IM, Inner-membrane.

## Supporting information

List of genes with unmodified expression

## Acknowledgment

The authors would like to thank Dre Wai from the Laboratory for Molecular Infection Medicine Sweden (MIMS) at Umeå University for bacterial strains. This work was supported by the Natural Sciences and Engineering Research Council of Canada (NSERC; http://www.nserc-crsng.gc.ca/index_eng.asp) Discovery grant number RGPIN-2017-05322 to MD. AM-D received financial support from the NSERC scholarship program (BESC D3 – 558624 – 2021). JP-F received financial support from the FRQNT Scholarship Program, the NSERC’s Canada Graduate Scholarships Master’s program (CGS M) and the J.A. DeSève Scholarship from the Graduate and Postdoctoral Studies of the University of Montreal (ESP). AM-D and MD received financial support from the RAQ (Ressources Aquatiques Québec), an inter-institutional group supported financially by the Fonds de recherche du Québec – Nature et technologies (FRQNT) (Programme regroupements stratégiques). GBW received financial support from a FRQNT postdoctoral fellowship and is financially supported by YVB through a Canada 150 Research Chair in Bacterial Cell Biology and Project Grant PJT-169053 from the Canadian Institutes of Health Research (CIHR).

## Authors contribution

Conceptualization: A.M.D., M.D.

Methodology: A.M.D., G.B.W., A.T.V., J.P.F., F.M.

Validation: All authors.

Formal analysis: A.M.D., G.B.W., A.T.V., J.P.F., M.D.

Investigation: A.M.D., G.B.W., A.T.V., J.P.F., M.D.

Resources: A.T.V., Y.V.B., M.D.

Data Curation: A.M.D., G.B.W., A.T.V., J.P.F., M.D.

Writing – Original Draft: A.M.D., G.B.W., M.D.

Writing – Review & Editing: All authors.

Visualization: A.M.D., G.B.W., M.D.

Supervision: Y.V.B., M.D.

Project administration: A.M.D., M.D.

Funding acquisition: M.D.

