## Supplementary material for "The *vprAB-ompV-virK* operon of *Vibrio cholerae* senses antimicrobial peptides and activates the expression of multiple resistance systems"

**- Supplementary materials –**

**Annabelle Mathieu-Denoncourt^1,2^, Gregory B. Whitfield^1,2^, Antony T. Vincent^3,4^, Julien Pauzé-Foixet^1,2^, Feriel Mahieddine^1,2^, Yves V. Brun^1,2^ and Marylise Duperthuy^1,2,^***

**^1^**Département de Microbiologie, infectiologie et immunologie, Faculté de médecine, Université de Montréal, Montréal, H3T 1J4, Québec, Canada.

**^2^**Centre d’Innovation Biomédicale, Faculté de médecine, Université de Montréal, Montréal, H3T 1J4, Québec, Canada.

**^3^**Département des sciences animales, Faculté des sciences de l'agriculture et de l'alimentation, Université Laval, Québec, G1V 0A6, Québec, Canada.

**^4^**Institut de biologie Intégrative et des systèmes, Université Laval, Québec, G1V 0A, Québec, Canada

*Corresponding author: Marylise Duperthuy, Université de Montréal, C.P. 6128, succursale Centre-ville, Montréal, QC, H3C 3J7, tel : 514 343-6111, fax:  514 343-5701,

1. **Supplementary figures**

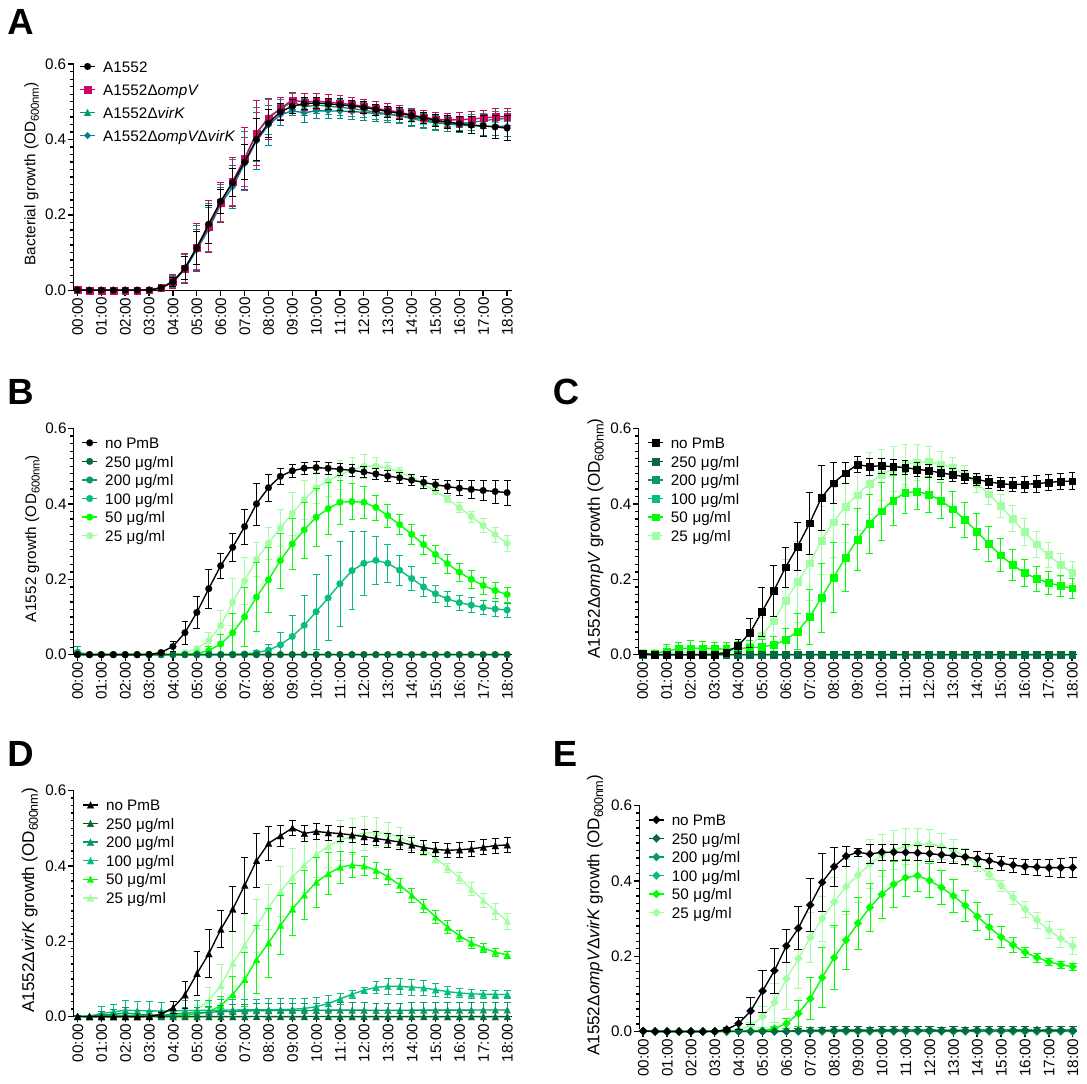

**Figure S1. The loss of *ompV* or *virK* has no effect on bacterial growth, but the loss of *ompV* and *virK* increases the susceptibility to polymyxin B.** **A)** Growth curves of A1552, A1552Δ*ompV,* A1552Δ*virK,* and A1552Δ*ompV*Δ*virK* in LB at 37°C in P96 plates, determined by measuring the optical density at 600 nm (OD_600nm_) over time. The growth curves were also determined in the presence of different concentrations of polymyxin B (PmB) for **B)** A1552**, C)** A1552Δ*ompV,* **D)** A1552Δ*virK* and **E)** A1552Δ*ompV*Δ*virK* in LB at 37°C*.* Results are presented as mean ± SD and were obtained from 3 independent experiments.

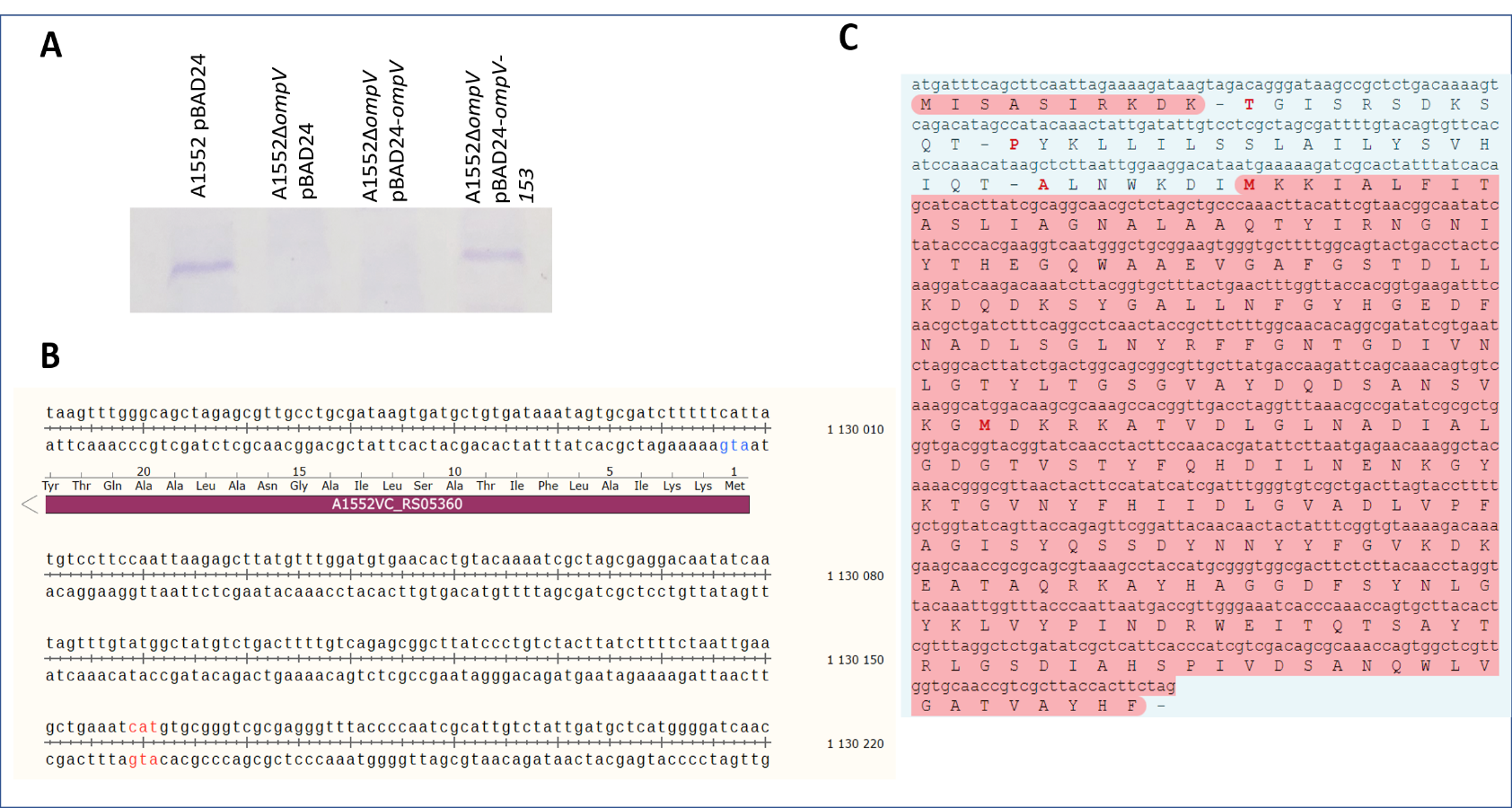

**Figure S2. Confirmation of *ompV* complementation using pBAD24 by the presence of OmpV in the membrane vesicles.** As OmpV was first observed in the membrane vesicles from cultures supplemented with PmB, vesicles were isolated from A1552 pBAD24, A1552Δ*ompV* pBAD24, A1552Δ*ompV* pBAD24-*ompV* and A1552Δ*ompV* pBAD24-*ompV*-153 cell-free supernatants grown for 16h in the presence of 3 µg/ml of polymyxin B. The vesicular proteins were migrated on a 13% SDS polyacrylamide gel stained with Coomassie blue. The results are representative of 3 independent experiments.

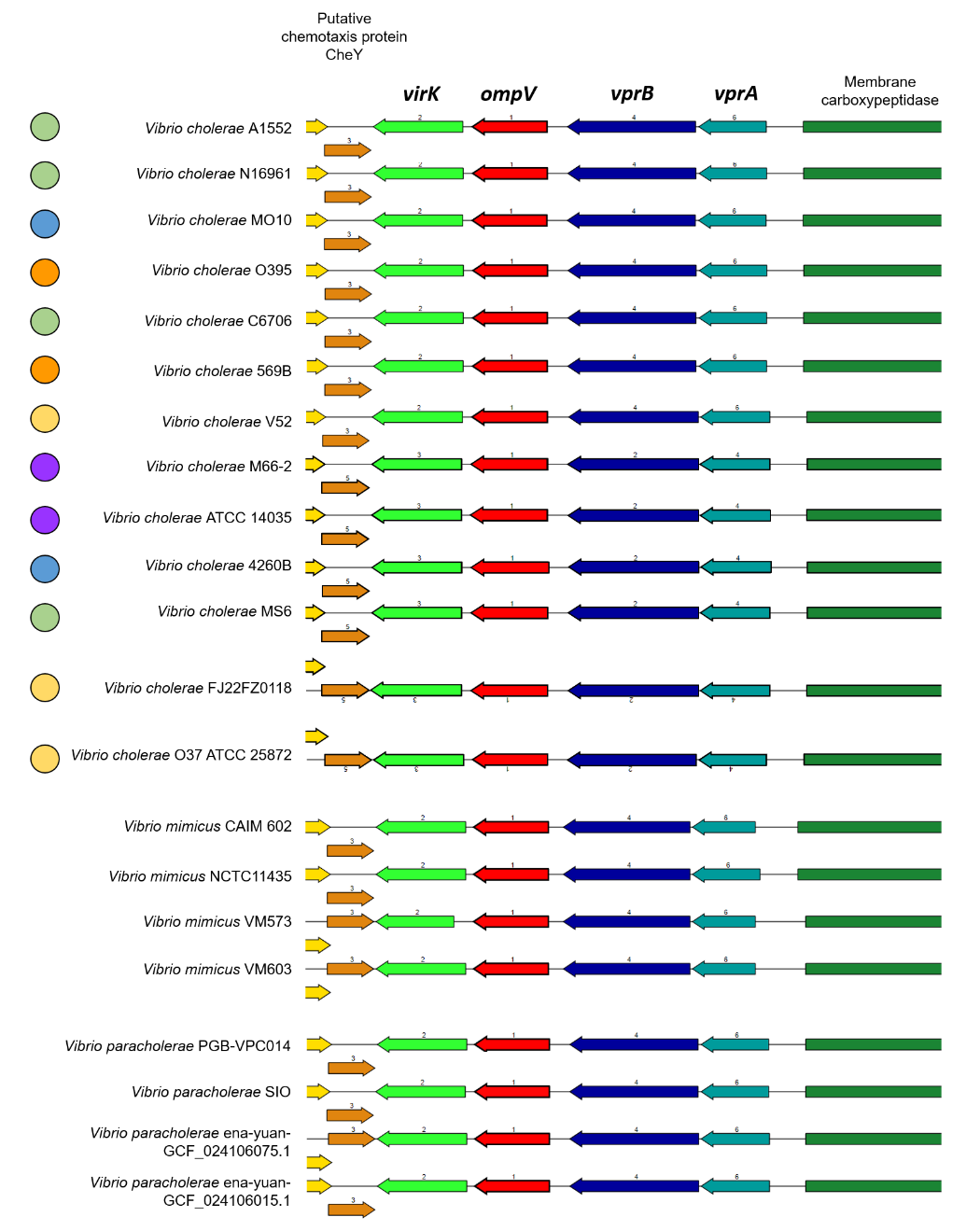

**Figure S3. The *vprAB-ompV-virK* operon is conserved amongst *Vibrio***. The *ompV* genomic region of *V. cholerae* A1552 was aligned with other *Vibrio* genomes with the Compare Region Viewer service of PATRIC (<https://www.patricbrc.org>) (1, 2). The *vprA-vprB-ompV-virK* operon is conserved amongst *V. cholerae*, including O1 El Tor (green dots) and Classical (orange dots), O139 (blue dots), and environmental strains (yellow dots). The locus is also conserved in *V. mimicus* and *V. paracholerae*. Purple dots, pre-pandemic O1 strains.

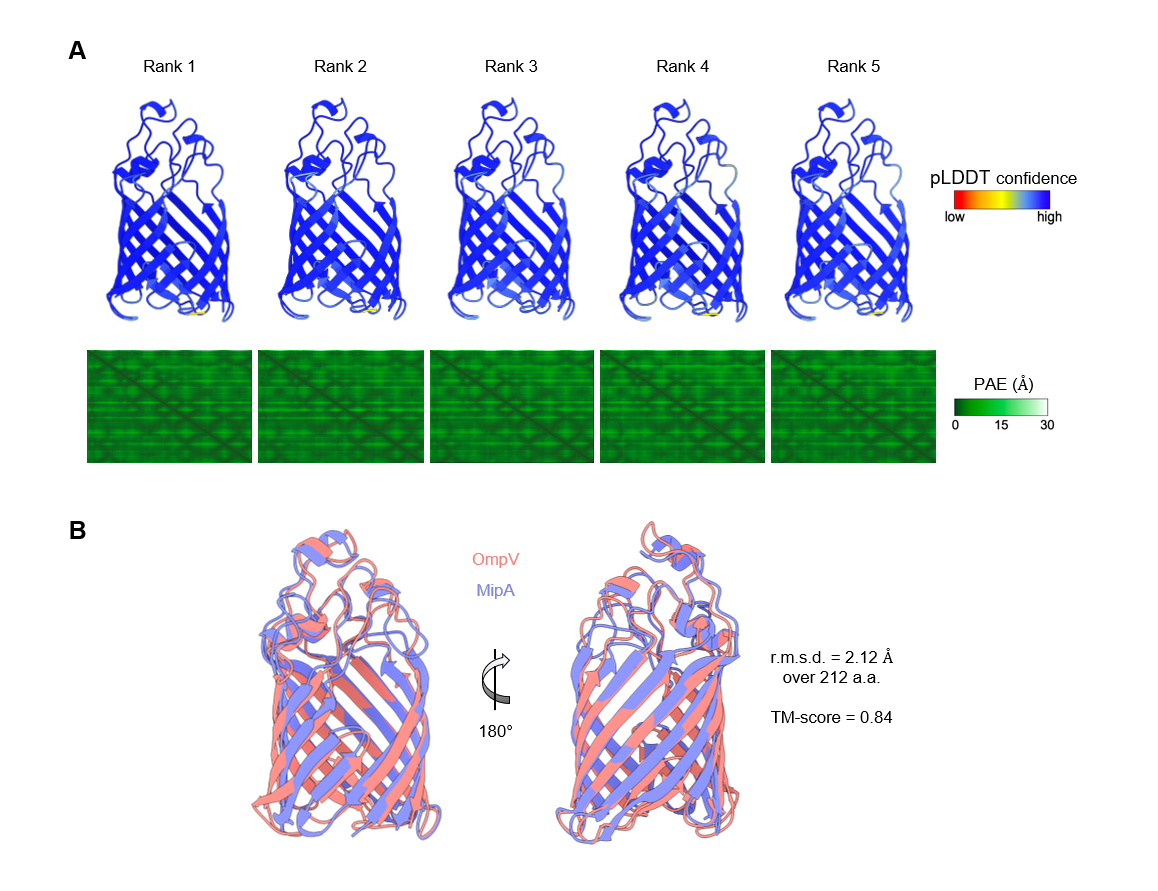

**Figure S4. Model confidences for the Alphafold2-predicted structure of OmpV, and alignment with *P. aeruginosa* MipA. A)** OmpV structures, with the predicted signal sequence (residues 1-19) removed, predicted by AlphaFold2 as implemented within ColabFold. Structures are aligned and shown from the same orientation. Structures are ranked according to the predicted template modeling (pTM) score and are colored according to the predicted local distance difference test (pLDDT) score, which indicates per-residue model confidence. The predicted aligned error (PAE) scores indicate positional error in angstroms for a given pair of residues across the entire protein chain. **B)** Alignment of the predicted structure of OmpV (pink) and the predicted structure of MipA (AFDB AF-A0A424XHT2-F1; purple). The root mean square deviation (r.m.s.d.) and template modeling (TM) score for the alignment are as indicated.

2. **Supplementary tables**

**Table SI. Genes with a modified expression in the presence of subinhibitory concentration of polymyxin B in *V. cholerae* A1552**

| **Locus tag** | **ID** | **Chromosome** | **Gene** | **baseMean** | **Log_2_(Fold Change)** | **lfcSE** | **stat** | **p-value** | **p-adj** | **Description** |
| --- | --- | --- | --- | --- | --- | --- | --- | --- | --- | --- |
| VC_RS08785 | VC_1820 | I |  | 140,6 | 4,19792 | 0,60123 | 6,98217 | 2,907E-12 | 4,231E-10 | PTS sugar transporter subunit IIA |
| VC_RS13475 | VC_A0013 | II |  | 2232,1 | 3,00260 | 0,75498 | 3,97706 | 6,977E-05 | 0,002134 | glycogen/starch/alpha-glucan phosphorylase |
| VC_RS18190 | VC_A1028 | II | *lamB* | 2257,8 | 2,87610 | 0,79625 | 3,61207 | 0,000304 | 0,007086 | maltoporin LamB |
| VC_RS06535 | VC_1343 | I |  | 528,0 | 2,48660 | 0,80179 | 3,10133 | 0,001927 | 0,030749 | tripeptidase T |
| VC_RS10735 | VC_2221 | I |  | 803,4 | 2,26056 | 0,47279 | 4,78134 | 1,741E-06 | 9,458E-05 | DUF3149 domain-containing protein |
| VC_RS15865 | VC_A0511 | II | *nrdD* | 3627,5 | 2,15803 | 0,48179 | 4,47915 | 7,494E-06 | 0,000345 | anaerobic ribonucleoside-triphosphate reductase |
| VC_RS13480 | VC_A0014 | II | *malQ* | 614,5 | 2,10599 | 0,54283 | 3,87964 | 0,000105 | 0,002906 | 4-alpha-glucanotransferase |
| VC_RS14210 | VC_A0180 | II | *pepT* | 867,5 | 2,03576 | 0,66881 | 3,04387 | 0,002336 | 0,034549 | peptidase T |
| VC_RS06410 | VC_1315 | I |  | 662,7 | 2,01775 | 0,22825 | 8,84023 | 9,552E-19 | 6,952E-16 | ATP-binding protein |
| VC_RS18380 | VC_A1069 | II |  | 3498,4 | 1,97247 | 0,66251 | 2,97726 | 0,002908 | 0,040706 | methyl-accepting chemotaxis protein |
| VC_RS15870 | VC_A0512 | II | *nrdG* | 291,3 | 1,90977 | 0,45804 | 4,16942 | 3,054E-05 | 0,001079 | anaerobic ribonucleoside-triphosphate reductase-activating protein |
| VC_RS08815 | VC_1826 | I |  | 236,7 | 1,88882 | 0,36792 | 5,13381 | 2,839E-07 | 1,913E-05 | fructose-specific PTS transporter subunit EIIC |
| VC_RS03300 | VC_0651 | I |  | 1426,1 | 1,77299 | 0,53870 | 3,29121 | 0,000998 | 0,017655 | U32 family peptidase |
| VC_RS17455 | VC_A0867 | II | *ompW* | 5806,6 | 1,75364 | 0,42123 | 4,16315 | 3,139E-05 | 0,001098 | outer membrane protein OmpW |
| VC_RS09030 | VC_1871 | I |  | 2915,1 | 1,69632 | 0,42330 | 4,00742 | 6,139E-05 | 0,001909 | YfbU family protein |
| VC_RS17815 | VC_A0944 | II | *malF* | 3392,6 | 1,67616 | 0,55631 | 3,01301 | 0,002587 | 0,036913 | maltose ABC transporter permease MalF |
| VC_RS06405 | VC_1314 | I |  | 601,8 | 1,66118 | 0,37581 | 4,42030 | 9,857E-06 | 0,000432 | SLC13 family permease |
| VC_RS10825 | VC_2240 | I |  | 3471,3 | 1,65187 | 0,29641 | 5,57299 | 2,504E-08 | 2,223E-06 | phenolic acid decarboxylase |
| VC_RS17215 | VC_A0811 | II | *gbpA* | 3157,4 | 1,64226 | 0,22179 | 7,40451 | 1,316E-13 | 2,994E-11 | N-acetylglucosamine-binding protein GbpA |
| VC_RS06430 | VC_1319 | I | *carS* | 28015,7 | 1,63536 | 0,16027 | 10,20401 | 1,903E-24 | 6,923E-21 | histidine kinase CarS |
| VC_RS17825 | VC_A0946 | II | *malK* | 4996,4 | 1,62882 | 0,50496 | 3,22563 | 0,001257 | 0,021374 | maltose/maltodextrin ABC transporter ATP-binding protein MalK |
| VC_RS03305 | VC_0652 | I |  | 1289,8 | 1,56734 | 0,34894 | 4,49169 | 7,066E-06 | 0,000330 | peptidase U32 family protein |
| VC_RS06420 | VC_1317 | I |  | 27657,0 | 1,56322 | 0,17491 | 8,93716 | 3,993E-19 | 3,633E-16 | VirK/YbjX family protein |
| VC_RS14180 | VC_A0174 | II |  | 14294,1 | 1,53512 | 0,18982 | 8,08712 | 6,109E-16 | 2,779E-13 | DUF58 domain-containing protein |
| VC_RS16560 | VC_A0665 | II | *dcuC* | 2073,0 | 1,52317 | 0,50481 | 3,01729 | 0,002550 | 0,036684 | anaerobic C4-dicarboxylate transporter DcuC |
| VC_RS14385 | VC_A0219 | II | *hlyA* | 9214,3 | 1,51016 | 0,28144 | 5,36584 | 8,057E-08 | 6,108E-06 | cytolysin VC_C |
| VC_RS08790 | VC_1821 | I |  | 644,6 | 1,49140 | 0,47012 | 3,17239 | 0,001512 | 0,024895 | fructose-specific PTS transporter subunit EIIC |
| VC_RS06415 | VC_1316 | I |  | 767,5 | 1,47412 | 0,20071 | 7,34448 | 2,066E-13 | 4,176E-11 | response regulator |
| VC_RS05470 | VC_1115 | I | *bioD* | 988,9 | 1,45972 | 0,46411 | 3,14523 | 0,001660 | 0,027081 | dethiobiotin synthase |
| VC_RS18000 | VC_A0985 | II |  | 1615,7 | 1,44897 | 0,25772 | 5,62228 | 1,885E-08 | 1,751E-06 | FAD-binding and (Fe-S)-binding domain-containing protein |
| VC_RS17995 | VC_A0984 | II | *lldD* | 470,5 | 1,41669 | 0,18884 | 7,50217 | 6,277E-14 | 1,632E-11 | FMN-dependent L-lactate dehydrogenase LldD |
| VC_RS12800 | VC_2657 | I |  | 3804,3 | 1,35621 | 0,16963 | 7,99528 | 1,293E-15 | 5,227E-13 | succinate dehydrogenase/fumarate reductase iron-sulfur subunit |
| VC_RS07565 | VC_1565 | I |  | 4417,2 | 1,34625 | 0,17317 | 7,77421 | 7,592E-15 | 2,763E-12 | TolC family protein |
| VC_RS07565 | VC_1565 | I |  | 4417,2 | 1,34625 | 0,17317 | 7,77421 | 7,592E-15 | 2,763E-12 | TolC family protein |
| VC_RS07575 | VC_1567 | I |  | 3595,9 | 1,34591 | 0,17888 | 7,52397 | 5,314E-14 | 1,487E-11 | ABC transporter permease |
| VC_RS16235 | VC_A0591 | II |  | 1521,9 | 1,34397 | 0,18954 | 7,09054 | 1,336E-12 | 2,066E-10 | extracellular solute-binding protein |
| VC_RS17990 | VC_A0983 | II |  | 762,7 | 1,34309 | 0,38799 | 3,46168 | 0,000537 | 0,010755 | L-lactate permease |
| VC_RS12795 | VC_2656 | I | *frdA* | 13524,3 | 1,34269 | 0,21082 | 6,36894 | 1,903E-10 | 2,309E-08 | fumarate reductase (quinol) flavoprotein subunit |
| VC_RS16230 | VC_A0590 | II |  | 2355,7 | 1,33569 | 0,18845 | 7,08781 | 1,363E-12 | 2,066E-10 | microcin C ABC transporter permease YejB |
| VC_RS07585 | VC_1570 | I |  | 500,7 | 1,32840 | 0,23406 | 5,67556 | 1,382E-08 | 1,360E-06 | cytochrome d ubiquinol oxidase subunit II |
| VC_RS16610 | VC_A0678 | II | *napA* | 23062,9 | 1,32012 | 0,31215 | 4,22910 | 2,346E-05 | 0,000880 | periplasmic nitrate reductase subunit alpha |
| VC_RS17915 | VC_A0966 | II |  | 480,7 | 1,31571 | 0,19337 | 6,80403 | 1,017E-11 | 1,424E-09 | hypothetical protein |
| VC_RS06435 | VC_1320 | I | *carR* | 9349,9 | 1,30716 | 0,14285 | 9,15076 | 5,654E-20 | 6,858E-17 | response regulator transcription factor CarR |
| VC_RS07315 | Pseudo | I |  | 6717,4 | 1,30582 | 0,36609 | 3,56691 | 0,000361 | 0,008164 | formate dehydrogenase subunit alpha |
| VC_RS07320 | VC_1514 | I |  | 461,7 | 1,29858 | 0,38933 | 3,33538 | 0,000852 | 0,015815 | twin-arginine translocation signal domain-containing protein |
| VC_RS06000 | VC_1224 | I | *relV* | 6672,5 | 1,28291 | 0,17986 | 7,13270 | 9,842E-13 | 1,628E-10 | (p)ppGpp synthetase RelV |
| VC_RS00720 | VC_0157 | I |  | 2906,1 | 1,27986 | 0,20685 | 6,18733 | 6,119E-10 | 6,959E-08 | S8 family peptidase |
| VC_RS07570 | VC_1566 | I |  | 2916,5 | 1,27912 | 0,14868 | 8,60340 | 7,739E-18 | 4,694E-15 | ABC transporter permease |
| VC_RS14170 | VC_A0172 | II |  | 9309,8 | 1,27895 | 0,13495 | 9,47716 | 2,613E-21 | 4,754E-18 | VWA domain-containing protein |
| VC_RS14185 | VC_A0175 | II |  | 25351,2 | 1,24723 | 0,24479 | 5,09504 | 3,487E-07 | 2,266E-05 | MoxR family ATPase |
| VC_RS02845 | VC_0554 | I |  | 22391,4 | 1,24662 | 0,16219 | 7,68601 | 1,518E-14 | 4,603E-12 | pitrilysin family protein |
| VC_RS07325 | VC_1515 | I |  | 1500,3 | 1,22933 | 0,37222 | 3,30272 | 0,000958 | 0,017165 | molecular chaperone |
| VC_RS14165 | VC_A0171 | II |  | 12157,0 | 1,22505 | 0,16764 | 7,30768 | 2,718E-13 | 4,945E-11 | VWA domain-containing protein |
| VC_RS09005 | VC_1866 | I | *pflB* | 180771,1 | 1,18701 | 0,31287 | 3,79400 | 0,000148 | 0,003881 | formate C-acetyltransferase |
| VC_RS17420 | VC_A0860 | II |  | 737,2 | 1,18483 | 0,39158 | 3,02580 | 0,002480 | 0,036095 | alpha-amylase |
| VC_RS07310 | VC_1512 | I | *fdh3B* | 254,2 | 1,17020 | 0,39613 | 2,95409 | 0,003136 | 0,043158 | formate dehydrogenase FDH3 subunit beta |
| VC_RS06425 | VC_1318 | I |  | 366207,5 | 1,14563 | 0,13781 | 8,31295 | 9,335E-17 | 4,853E-14 | MipA/OmpV family protein |
| VC_RS16220 | VC_A0588 | II |  | 2155,1 | 1,14063 | 0,22918 | 4,97706 | 6,456E-07 | 3,851E-05 | ABC transporter ATP-binding protein |
| VC_RS07680 | VC_1590 | I | *alsS* | 1342,5 | 1,13834 | 0,20724 | 5,49295 | 3,953E-08 | 3,197E-06 | acetolactate synthase AlsS |
| VC_RS17100 | VC_A0784 | II |  | 1954,5 | 1,13265 | 0,36719 | 3,08462 | 0,002038 | 0,031561 | trans-2-enoyl-CoA reductase family protein |
| VC_RS07955 | VC_1642 | I |  | 487,5 | 1,11933 | 0,17966 | 6,23042 | 4,652E-10 | 5,461E-08 | ribosome recycling factor family protein |
| VC_RS02900 | VC_0566 | I |  | 51950,6 | 1,11886 | 0,19417 | 5,76239 | 8,293E-09 | 8,876E-07 | DegQ family serine endoprotease |
| VC_RS13005 | VC_2701 | I |  | 5004,8 | 1,11702 | 0,15124 | 7,38558 | 1,518E-13 | 3,249E-11 | protein-disulfide reductase DsbD |
| VC_RS06100 | VC_1249 | I |  | 519,4 | 1,11171 | 0,26173 | 4,24762 | 2,161E-05 | 0,000828 | glycine cleavage system protein R |
| VC_RS07560 | Pseudo | I |  | 7164,7 | 1,11129 | 0,21857 | 5,08443 | 3,687E-07 | 2,313E-05 | efflux RND transporter periplasmic adaptor subunit |
| VC_RS07370 | VC_1524 | I |  | 1364,6 | 1,10703 | 0,35885 | 3,08492 | 0,002036 | 0,031561 | ABC transporter permease |
| VC_RS12810 | VC_2659 | I | *frdD* | 2270,4 | 1,09614 | 0,21276 | 5,15191 | 2,579E-07 | 1,781E-05 | fumarate reductase subunit FrdD |
| VC_RS12805 | VC_2658 | I | *frdC* | 2066,3 | 1,09140 | 0,21559 | 5,06238 | 4,141E-07 | 2,511E-05 | fumarate reductase subunit FrdC |
| VC_RS14175 | VC_A0173 | II |  | 6681,8 | 1,08957 | 0,23303 | 4,67569 | 2,930E-06 | 0,000152 | DUF4381 family protein |
| VC_RS07580 | VC_1568 | I |  | 3028,9 | 1,08323 | 0,14983 | 7,22964 | 4,843E-13 | 8,392E-11 | ABC transporter ATP-binding protein |
| VC_RS03110 | VC_0614 | I |  | 159,7 | 1,07540 | 0,25360 | 4,24049 | 2,230E-05 | 0,000845 | N-acetylglucosamine kinase |
| VC_RS03400 | VC_0673 | I |  | 9361,3 | 1,05652 | 0,14441 | 7,31629 | 2,549E-13 | 4,882E-11 | sulfite exporter TauE/SafE family protein |
| VC_RS01365 | VC_0281 | I |  | 509,6 | 1,03521 | 0,18324 | 5,64960 | 1,608E-08 | 1,540E-06 | lysine decarboxylase CadA |
| VC_RS13625 | VC_A0047 | II |  | 12210,5 | 1,03062 | 0,15180 | 6,78946 | 1,126E-11 | 1,517E-09 | HlyD family secretion protein |
| VC_RS14160 | Pseudo | II |  | 14584,2 | 1,01600 | 0,15811 | 6,42581 | 1,312E-10 | 1,646E-08 | BatD family protein |
| VC_RS13630 | VC_A0048 | II |  | 11606,8 | 1,00745 | 0,18150 | 5,55080 | 2,844E-08 | 2,412E-06 | DUF3302 domain-containing protein |
| VC_RS02530 | VC_0496 | I |  | 413,3 | 1,00185 | 0,19701 | 5,08520 | 3,672E-07 | 2,313E-05 | inovirus-type Gp2 protein |
| VC_RS08410 | NA | I |  | 1810,4 | 0,99817 | 0,22077 | 4,52131 | 6,146E-06 | 0,000290 | hypothetical protein |
| VC_RS08100 | VC_1674 | I |  | 475,1 | 0,99045 | 0,17917 | 5,52798 | 3,239E-08 | 2,679E-06 | efflux RND transporter periplasmic adaptor subunit |
| VC_RS17895 | VC_A0962 | II |  | 2927,0 | 0,98250 | 0,26128 | 3,76029 | 0,000170 | 0,004308 | DUF3541 domain-containing protein |
| VC_RS17845 | VC_A0951 | II |  | 5509,4 | 0,97409 | 0,20052 | 4,85776 | 1,187E-06 | 6,647E-05 | heavy metal-binding domain-containing protein |
| VC_RS07185 | VC_1485 | I |  | 53810,8 | 0,97128 | 0,13016 | 7,46230 | 8,502E-14 | 2,063E-11 | DUF3466 family protein |
| VC_RS01780 | VC_0353 | I |  | 18519,2 | 0,96418 | 0,16760 | 5,75292 | 8,772E-09 | 9,120E-07 | hypothetical protein |
| VC_RS17850 | VC_A0952 | II | *vpsT* | 4673,9 | 0,94257 | 0,16162 | 5,83202 | 5,476E-09 | 6,039E-07 | LuxR family transcriptional regulator VpsT |
| VC_RS16225 | VC_A0589 | II |  | 592,1 | 0,94196 | 0,21918 | 4,29774 | 1,725E-05 | 0,000690 | ABC transporter permease |
| VC_RS02840 | VC_0553 | I |  | 1704,3 | 0,94084 | 0,16470 | 5,71243 | 1,114E-08 | 1,126E-06 | YqaA family protein |
| VC_RS17910 | VC_A0965 | II |  | 2216,8 | 0,93821 | 0,20297 | 4,62252 | 3,791E-06 | 0,000190 | diguanylate cyclase |
| VC_RS02960 | VC_0580 | I |  | 6156,6 | 0,93548 | 0,23315 | 4,01238 | 6,011E-05 | 0,001902 | YraN family protein |
| VC_RS08415 | VC_1743 | I |  | 1970,3 | 0,92300 | 0,23818 | 3,87524 | 0,000107 | 0,002937 | 1-aminocyclopropane-1-carboxylate deaminase/D-cysteine desulfhydrase |
| VC_RS09560 | VC_1987 | I |  | 2466,2 | 0,92103 | 0,17025 | 5,40995 | 6,304E-08 | 4,881E-06 | Slp family lipoprotein |
| VC_RS16600 | VC_A0676 | II | *napF* | 4748,7 | 0,91165 | 0,28679 | 3,17882 | 0,001479 | 0,024637 | ferredoxin-type protein NapF |
| VC_RS06785 | VC_1401 | I |  | 354,8 | 0,91088 | 0,19336 | 4,71083 | 2,467E-06 | 0,000132 | chemotaxis protein CheB |
| VC_RS00160 | VC_0035 | I |  | 6816,6 | 0,90157 | 0,21438 | 4,20543 | 2,606E-05 | 0,000958 | serine/threonine protein kinase |
| VC_RS07940 | VC_1639 | I |  | 1408,1 | 0,87043 | 0,15682 | 5,55038 | 2,850E-08 | 2,412E-06 | ATP-binding protein |
| VC_RS13925 | VC_A0116 | II | *tssH* | 1245,9 | 0,85989 | 0,23747 | 3,62101 | 0,000293 | 0,006890 | type VI secretion system ATPase TssH |
| VC_RS05695 | VC_1162 | I |  | 3387,7 | 0,85827 | 0,16663 | 5,15084 | 2,593E-07 | 1,781E-05 | ATP-dependent zinc protease |
| VC_RS07330 | VC_1516 | I |  | 4373,3 | 0,85556 | 0,22137 | 3,86493 | 0,000111 | 0,003006 | 4Fe-4S dicluster domain-containing protein |
| VC_RS13165 | Pseudo | I | *gspD* | 8830,9 | 0,85473 | 0,16734 | 5,10771 | 3,261E-07 | 2,157E-05 | type II secretion system secretin GspD |
| VC_RS13935 | VC_A0118 | II | *vasI* | 204,2 | 0,85349 | 0,22952 | 3,71864 | 0,000200 | 0,004958 | type VI secretion system-associated protein VasI |
| VC_RS11640 | VC_2417 | I | *recJ* | 5535,2 | 0,85075 | 0,15142 | 5,61862 | 1,925E-08 | 1,751E-06 | single-stranded-DNA-specific exonuclease RecJ |
| VC_RS09330 | VC_1936 | I |  | 1686,1 | 0,84397 | 0,23869 | 3,53588 | 0,000406 | 0,008887 | phosphatidate cytidylyltransferase |
| VC_RS11885 | VC_2467 | I | *rpoE* | 34812,1 | 0,84335 | 0,18310 | 4,60599 | 4,105E-06 | 0,000197 | RNA polymerase sigma factor RpoE |
| VC_RS05690 | VC_1161 | I |  | 5098,4 | 0,82166 | 0,19003 | 4,32379 | 1,534E-05 | 0,000627 | inactive transglutaminase family protein |
| VC_RS00775 | VC_0164 | I | *vexB* | 48888,6 | 0,81869 | 0,21808 | 3,75407 | 0,000174 | 0,004366 | multidrug efflux RND transporter permease subunit VexB |
| VC_RS00155 | VC_0034 | I |  | 44973,8 | 0,81597 | 0,15526 | 5,25544 | 1,477E-07 | 1,075E-05 | thiol:disulfide interchange protein DsbA/DsbL |
| VC_RS16615 | VC_A0679 | II |  | 6762,3 | 0,81425 | 0,27883 | 2,92023 | 0,003498 | 0,047108 | nitrate reductase cytochrome c-type subunit |
| VC_RS16350 | VC_A0619 | II |  | 574,2 | 0,81243 | 0,26989 | 3,01021 | 0,002611 | 0,037111 | VOC family protein |
| VC_RS13930 | VC_A0117 | II | *vasH* | 865,3 | 0,80614 | 0,20931 | 3,85149 | 0,000117 | 0,003141 | sigma-54 dependent T6SS transcriptional regulator VasH |
| VC_RS07630 | VC_1579 | I | *almE* | 80594,7 | 0,79848 | 0,23056 | 3,46315 | 0,000534 | 0,010755 | lipid A modification system glycine--protein ligase AlmE |
| VC_RS13170 | VC_2734 | I | *gspC* | 4056,3 | 0,79678 | 0,15721 | 5,06833 | 4,013E-07 | 2,475E-05 | type II secretion system protein GspC |
| VC_RS03560 | VC_0708 | I |  | 29440,5 | 0,79380 | 0,24522 | 3,23714 | 0,001207 | 0,020633 | outer membrane protein assembly factor BamD |
| VC_RS09325 | VC_1935 | I |  | 2252,4 | 0,78988 | 0,21373 | 3,69567 | 0,000219 | 0,005320 | CDP-alcohol phosphatidyltransferase family protein |
| VC_RS09335 | VC_1937 | I |  | 576,6 | 0,78781 | 0,18960 | 4,15508 | 3,252E-05 | 0,001127 | lysophospholipid acyltransferase family protein |
| VC_RS10400 | VC_2164 | I |  | 11791,8 | 0,78392 | 0,17576 | 4,46016 | 8,190E-06 | 0,000373 | M48 family metallopeptidase |
| VC_RS14550 | VC_A0255 | II |  | 244,5 | 0,78388 | 0,22121 | 3,54370 | 0,000395 | 0,008755 | DUF2861 family protein |
| VC_RS13160 | VC_2732 | I | *gspE* | 4421,5 | 0,78190 | 0,20105 | 3,88913 | 0,000101 | 0,002838 | type II secretion system ATPase GspE |
| VC_RS13490 | VC_A0017 | II | *hcp-2* | 795,6 | 0,78165 | 0,16939 | 4,61459 | 3,939E-06 | 0,000191 | type VI secretion system effector Hcp-2 |
| VC_RS12725 | VC_2638 | I |  | 632,1 | 0,77587 | 0,18254 | 4,25044 | 2,134E-05 | 0,000828 | dihydrolipoyl dehydrogenase |
| VC_RS00125 | VC_0028 | I | *ilvD* | 2224,5 | 0,77260 | 0,21008 | 3,67770 | 0,000235 | 0,005672 | dihydroxy-acid dehydratase |
| VC_RS13905 | VC_A0112 | II | *tagH* | 726,0 | 0,77012 | 0,26162 | 2,94365 | 0,003244 | 0,044212 | type VI secretion system-associated FHA domain protein TagH |
| VC_RS11645 | VC_2418 | I | *dsbC* | 7257,7 | 0,76804 | 0,14145 | 5,42986 | 5,640E-08 | 4,462E-06 | bifunctional protein-disulfide isomerase/oxidoreductase DsbC |
| VC_RS11650 | VC_2419 | I | *xerD* | 3359,0 | 0,76627 | 0,18328 | 4,18088 | 2,904E-05 | 0,001046 | site-specific tyrosine recombinase XerD |
| VC_RS11745 | VC_2438 | I | *glnE* | 2762,1 | 0,76107 | 0,20027 | 3,80015 | 0,000145 | 0,003813 | bifunctional (glutamate-ammonia ligase)-adenylyl-L-tyrosine phosphorylase/(glutamate-ammonia-ligase) adenylyltransferase |
| VC_RS13135 | VC_2727 | I | *gspJ* | 444,1 | 0,75695 | 0,20136 | 3,75917 | 0,000170 | 0,004308 | type II secretion system minor pseudopilin GspJ |
| VC_RS02850 | VC_0556 | I | *gshA* | 17579,4 | 0,75474 | 0,19533 | 3,86402 | 0,000112 | 0,003006 | glutamate--cysteine ligase |
| VC_RS07935 | VC_1638 | I |  | 1194,1 | 0,75383 | 0,17136 | 4,39922 | 1,086E-05 | 0,000471 | response regulator transcription factor |
| VC_RS15990 | VC_A0538 | II |  | 1326,3 | 0,74654 | 0,21774 | 3,42860 | 0,000607 | 0,011559 | cytochrome b |
| VC_RS08220 | VC_1698 | I |  | 1225,2 | 0,74627 | 0,25352 | 2,94363 | 0,003244 | 0,044212 | YdcF family protein |
| VC_RS11880 | VC_2466 | I |  | 17110,5 | 0,74381 | 0,21167 | 3,51394 | 0,000442 | 0,009341 | sigma-E factor negative regulatory protein |
| VC_RS10240 | NA | I |  | 216,4 | 0,74277 | 0,24428 | 3,04063 | 0,002361 | 0,034782 | hypothetical protein |
| VC_RS07685 | VC_1591 | I |  | 2014,7 | 0,73770 | 0,15022 | 4,91066 | 9,077E-07 | 5,328E-05 | SDR family oxidoreductase |
| VC_RS07620 | VC_1577 | I | *almG* | 21221,4 | 0,73610 | 0,18248 | 4,03383 | 5,487E-05 | 0,001752 | glycine--lipid A transferase AlmG |
| VC_RS07625 | VC_1578 | I | *almF* | 6824,5 | 0,73464 | 0,24052 | 3,05441 | 0,002255 | 0,034167 | lipid A modification system glycine carrier protein AlmF |
| VC_RS11805 | VC_2451 | I | *relA* | 9406,1 | 0,71462 | 0,14635 | 4,88298 | 1,045E-06 | 6,036E-05 | GTP diphosphokinase |
| VC_RS13120 | VC_2724 | I | *gspM* | 1354,6 | 0,71337 | 0,16420 | 4,34443 | 1,396E-05 | 0,000584 | type II secretion system protein GspM |
| VC_RS02245 | VC_0445 | I | *surA* | 11200,1 | 0,71286 | 0,18909 | 3,77004 | 0,000163 | 0,004243 | peptidylprolyl isomerase SurA |
| VC_RS10410 | VC_2166 | I | *wrbA* | 6015,7 | 0,71106 | 0,13464 | 5,28129 | 1,283E-07 | 9,527E-06 | NAD(P)H:quinone oxidoreductase |
| VC_RS10875 | VC_2251 | I |  | 40889,1 | 0,70815 | 0,17664 | 4,00904 | 6,097E-05 | 0,001909 | OmpH family outer membrane protein |
| VC_RS08225 | VC_1699 | I |  | 673,4 | 0,70557 | 0,20441 | 3,45166 | 0,000557 | 0,011019 | hypothetical protein |
| VC_RS16045 | VC_A0551 | II |  | 648,1 | 0,70525 | 0,19784 | 3,56469 | 0,000364 | 0,008183 | hypothetical protein |
| VC_RS08105 | VC_1675 | I |  | 635,5 | 0,69510 | 0,19888 | 3,49508 | 0,000474 | 0,009799 | efflux RND transporter periplasmic adaptor subunit |
| VC_RS16215 | VC_A0587 | II |  | 380,9 | 0,69279 | 0,19616 | 3,53180 | 0,000413 | 0,008887 | DNA-binding protein |
| VC_RS17360 | VC_A0846 | II |  | 2445,9 | 0,68773 | 0,21245 | 3,23705 | 0,001208 | 0,020633 | LysE family translocator |
| VC_RS07545 | VC_1560 | I | *katG* | 1677,7 | 0,68643 | 0,16928 | 4,05496 | 5,014E-05 | 0,001627 | catalase/peroxidase HPI |
| VC_RS08095 | VC_1673 | I | *vexK* | 1517,9 | 0,68385 | 0,19672 | 3,47622 | 0,000509 | 0,010338 | efflux RND transporter permease subunit VexK |
| VC_RS13915 | VC_A0114 | II | *tssK* | 804,8 | 0,68114 | 0,21446 | 3,17612 | 0,001493 | 0,024689 | type VI secretion system baseplate subunit TssK |
| VC_RS11870 | VC_2464 | I |  | 3601,5 | 0,67757 | 0,13930 | 4,86413 | 1,150E-06 | 6,537E-05 | SoxR reducing system RseC family protein |
| VC_RS07790 | VC_1608 | I |  | 1071,5 | 0,67529 | 0,19193 | 3,51847 | 0,000434 | 0,009237 | ABC transporter permease |
| VC_RS13155 | VC_2731 | I | *gspF* | 3965,1 | 0,67307 | 0,17881 | 3,76419 | 0,000167 | 0,004282 | type II secretion system inner membrane protein GspF |
| VC_RS13125 | VC_2725 | I | *gspL* | 3048,6 | 0,66247 | 0,14261 | 4,64540 | 3,394E-06 | 0,000174 | type II secretion system protein GspL |
| VC_RS02955 | VC_0579 | I |  | 14956,0 | 0,65941 | 0,14269 | 4,62118 | 3,816E-06 | 0,000190 | phosphoheptose isomerase |
| VC_RS13505 | VC_A0020 | II | *vasX* | 2005,7 | 0,65224 | 0,19014 | 3,43036 | 0,000603 | 0,011545 | type VI secretion system toxin VasX |
| VC_RS07360 | VC_1522 | I |  | 767,5 | 0,64991 | 0,21013 | 3,09283 | 0,001983 | 0,031330 | sigma-54 dependent transcriptional regulator |
| VC_RS10250 | VC_2130 | I | *fliI* | 1328,3 | 0,64879 | 0,15217 | 4,26351 | 2,012E-05 | 0,000796 | flagellar protein export ATPase FliI |
| VC_RS09495 | VC_1973 | I | *menB* | 6082,4 | 0,64632 | 0,16155 | 4,00073 | 6,315E-05 | 0,001947 | 1,4-dihydroxy-2-naphthoyl-CoA synthase |
| VC_RS11405 | VC_2367 | I |  | 955,9 | 0,64623 | 0,16436 | 3,93193 | 8,427E-05 | 0,002453 | DUF3293 domain-containing protein |
| VC_RS06110 | VC_1252 | I |  | 4124,5 | 0,64585 | 0,20898 | 3,09041 | 0,001999 | 0,031330 | CinA family nicotinamide mononucleotide deamidase-related protein |
| VC_RS00780 | VC_0165 | I | *vexA* | 29309,9 | 0,64426 | 0,19901 | 3,23733 | 0,001207 | 0,020633 | multidrug efflux RND transporter periplasmic adaptor subunit VexA |
| VC_RS07600 | VC_1573 | I |  | 590,2 | 0,63968 | 0,17234 | 3,71178 | 0,000206 | 0,005026 | class II fumarate hydratase |
| VC_RS00615 | VC_0132 | I |  | 3699,6 | 0,63586 | 0,14338 | 4,43467 | 9,222E-06 | 0,000409 | COG3650 family protein |
| VC_RS10265 | VC_2133 | I | *fliF* | 4928,4 | 0,62617 | 0,14518 | 4,31311 | 1,610E-05 | 0,000651 | flagellar basal-body MS-ring/collar protein FliF |
| VC_RS05140 | VC_1045 | I |  | 5392,9 | 0,62358 | 0,15961 | 3,90678 | 9,353E-05 | 0,002682 | sigma-70 family RNA polymerase sigma factor |
| VC_RS11875 | VC_2465 | I | *rseB* | 11688,1 | 0,62318 | 0,19480 | 3,19904 | 0,001379 | 0,023230 | sigma-E factor regulatory protein RseB |
| VC_RS11400 | VC_2366 | I |  | 1603,9 | 0,62277 | 0,17993 | 3,46113 | 0,000538 | 0,010755 | putative 4-hydroxy-4-methyl-2-oxoglutarate aldolase |
| VC_RS08315 | VC_1718 | I | *elyC* | 5089,3 | 0,61741 | 0,16394 | 3,76600 | 0,000166 | 0,004281 | envelope biogenesis factor ElyC |
| VC_RS10255 | VC_2131 | I | *fliH* | 1614,2 | 0,61405 | 0,14893 | 4,12300 | 3,740E-05 | 0,001265 | flagellar assembly protein FliH |
| VC_RS13130 | VC_2726 | I | *gspK* | 869,5 | 0,60858 | 0,17768 | 3,42520 | 0,000614 | 0,011644 | type II secretion system minor pseudopilin GspK |
| VC_RS09490 | VC_1972 | I | *menC* | 543,6 | 0,60811 | 0,20842 | 2,91772 | 0,003526 | 0,047108 | o-succinylbenzoate synthase |
| VC_RS09965 | VC_2068 | I | *flhF* | 4971,8 | 0,60224 | 0,14362 | 4,19319 | 2,751E-05 | 0,001001 | flagellar biosynthesis protein FlhF |
| VC_RS09175 | VC_1902 | I | *dsbB* | 2990,7 | 0,60159 | 0,14842 | 4,05322 | 5,052E-05 | 0,001627 | disulfide bond formation protein DsbB |
| VC_RS10245 | VC_2129 | I | *fliJ* | 502,2 | 0,59625 | 0,17540 | 3,39941 | 0,000675 | 0,012667 | flagellar export protein FliJ |
| VC_RS07950 | VC_1641 | I |  | 1937,3 | 0,58818 | 0,15120 | 3,89014 | 0,000100 | 0,002838 | EAL domain-containing protein |
| VC_RS01785 | VC_0354 | I | *fkpA* | 121853,4 | 0,58641 | 0,17031 | 3,44307 | 0,000575 | 0,011203 | FKBP-type peptidyl-prolyl cis-trans isomerase |
| VC_RS09525 | VC_1979 | I |  | 1902,8 | 0,58620 | 0,16251 | 3,60715 | 0,000310 | 0,007175 | anti-phage deoxyguanosine triphosphatase |
| VC_RS14555 | VC_A0256 | II | *qseB* | 323,0 | 0,56985 | 0,19618 | 2,90464 | 0,003677 | 0,048478 | response regulator transcription factor |
| VC_RS09970 | VC_2069 | I | *flhA* | 4597,3 | 0,56886 | 0,14447 | 3,93769 | 8,227E-05 | 0,002434 | flagellar biosynthesis protein FlhA |
| VC_RS02950 | VC_0578 | I |  | 4762,4 | 0,56817 | 0,14402 | 3,94518 | 7,974E-05 | 0,002398 | BON domain-containing protein |
| VC_RS00925 | VC_0187 | I |  | 3997,4 | 0,56796 | 0,13990 | 4,05983 | 4,911E-05 | 0,001610 | 23S rRNA (adenine(2030)-N(6))-methyltransferase RlmJ |
| VC_RS14565 | VC_A0258 | II |  | 745,6 | 0,56607 | 0,17111 | 3,30824 | 0,000939 | 0,016913 | cupin domain-containing protein |
| VC_RS08470 | VC_1756 | I | *vexC* | 4516,4 | 0,56021 | 0,17934 | 3,12364 | 0,001786 | 0,028762 | multidrug efflux RND transporter periplasmic adaptor subunit VexC |
| VC_RS16115 | VC_A0566 | II | *vxrB* | 9904,8 | 0,55840 | 0,14372 | 3,88528 | 0,000102 | 0,002861 | response regulator transcription factor VxrB |
| VC_RS08420 | VC_1744 | I |  | 647,6 | 0,55698 | 0,18810 | 2,96104 | 0,003066 | 0,042585 | YnjH family protein |
| VC_RS16240 | VC_A0592 | II |  | 633,1 | 0,54360 | 0,18154 | 2,99432 | 0,002751 | 0,038947 | NUDIX hydrolase |
| VC_RS10405 | VC_2165 | I | *arsC* | 2444,1 | 0,54351 | 0,15616 | 3,48053 | 0,000500 | 0,010231 | arsenate reductase (glutaredoxin) |
| VC_RS05135 | VC_1044 | I |  | 6576,1 | 0,54310 | 0,13902 | 3,90663 | 9,359E-05 | 0,002682 | DUF3379 domain-containing protein |
| VC_RS03405 | VC_0674 | I | *lgt* | 5288,7 | 0,54191 | 0,13756 | 3,93931 | 8,172E-05 | 0,002434 | prolipoprotein diacylglyceryl transferase |
| VC_RS09100 | VC_1886 | I | *mfd* | 6859,7 | 0,53993 | 0,15277 | 3,53435 | 0,000409 | 0,008887 | transcription-repair coupling factor |
| VC_RS16270 | VC_A0600 | II | *phnR* | 1139,6 | 0,53797 | 0,17008 | 3,16300 | 0,001562 | 0,025596 | phosphonate utilization transcriptional regulator PhnR |
| VC_RS13940 | VC_A0119 | II | *vasJ* | 559,0 | 0,52882 | 0,17905 | 2,95341 | 0,003143 | 0,043158 | TssA family type VI secretion system protein VasJ |
| VC_RS11740 | Pseudo | I | *hldE* | 8197,0 | 0,52839 | 0,14203 | 3,72032 | 0,000199 | 0,004958 | bifunctional D-glycero-beta-D-manno-heptose-7-phosphate kinase/D-glycero-beta-D-manno-heptose 1-phosphate adenylyltransferase HldE |
| VC_RS06850 | VC_1416 | I | *tssI* | 849,5 | 0,52810 | 0,16876 | 3,12927 | 0,001752 | 0,028343 | type VI secretion system tip protein TssI/VgrG |
| VC_RS08055 | VC_1665 | I |  | 887,8 | 0,52238 | 0,16948 | 3,08229 | 0,002054 | 0,031674 | ABC transporter permease |
| VC_RS05945 | VC_1214 | I | *uvrC* | 6172,5 | 0,51118 | 0,14424 | 3,54399 | 0,000394 | 0,008755 | excinuclease ABC subunit UvrC |
| VC_RS14155 | VC_A0167 | II |  | 15166,2 | 0,50749 | 0,13120 | 3,86818 | 0,000110 | 0,003000 | SIMPL domain-containing protein |
| VC_RS13945 | VC_A0120 | II | *tssM* | 2841,0 | 0,49891 | 0,13952 | 3,57587 | 0,000349 | 0,008038 | type VI secretion system membrane subunit TssM |
| VC_RS17020 | VC_A0765 | II | *ltaE* | 3544,2 | 0,49741 | 0,14190 | 3,50531 | 0,000456 | 0,009538 | low-specificity L-threonine aldolase |
| VC_RS04710 | VC_0950 | I | *mrdA* | 8967,5 | 0,48992 | 0,16161 | 3,03142 | 0,002434 | 0,035576 | penicillin-binding protein 2 |
| VC_RS09250 | VC_1918 | I | *ppiD* | 29624,6 | 0,48955 | 0,15954 | 3,06858 | 0,002151 | 0,032885 | peptidylprolyl isomerase |
| VC_RS02250 | VC_0446 | I | *lptD* | 41239,3 | 0,48831 | 0,13662 | 3,57426 | 0,000351 | 0,008038 | LPS assembly protein LptD |
| VC_RS11800 | VC_2450 | I | *mazG* | 3048,2 | 0,48486 | 0,14427 | 3,36081 | 0,000777 | 0,014503 | nucleoside triphosphate pyrophosphohydrolase |
| VC_RS09935 | VC_2062 | I |  | 9613,2 | 0,48469 | 0,13770 | 3,52002 | 0,000432 | 0,009237 | chemotaxis response regulator protein-glutamate methylesterase |
| VC_RS10930 | VC_2262 | I | *glnD* | 3899,6 | 0,48372 | 0,14532 | 3,32874 | 0,000872 | 0,016115 | bifunctional uridylyltransferase/uridylyl-removing protein GlnD |
| VC_RS13075 | VC_2716 | I |  | 7414,8 | 0,48244 | 0,13993 | 3,44775 | 0,000565 | 0,011119 | Tex family protein |
| VC_RS10235 | VC_2128 | I |  | 2518,2 | 0,47906 | 0,14405 | 3,32570 | 0,000882 | 0,016128 | flagellar hook-length control protein FliK |
| VC_RS10260 | VC_2132 | I | *fliG* | 2431,5 | 0,46342 | 0,15997 | 2,89688 | 0,003769 | 0,049336 | flagellar motor switch protein FliG |
| VC_RS08085 | VC_1671 | I |  | 3042,3 | 0,46163 | 0,15291 | 3,01897 | 0,002536 | 0,036684 | cystathionine beta-lyase |
| VC_RS04700 | VC_0948 | I | *rlpA* | 17911,7 | 0,45467 | 0,13668 | 3,32645 | 0,000880 | 0,016128 | septal ring lytic transglycosylase RlpA |
| VC_RS03815 | VC_0761 | I |  | 6081,3 | 0,44211 | 0,13435 | 3,29068 | 0,000999 | 0,017655 | YfgM family protein |
| VC_RS16120 | VC_A0567 | II |  | 10825,7 | 0,44174 | 0,13419 | 3,29197 | 0,000995 | 0,017655 | DUF2861 family protein |
| VC_RS12000 | VC_2492 | I | *leuC* | 2831,4 | 0,42916 | 0,14062 | 3,05194 | 0,002274 | 0,034189 | 3-isopropylmalate dehydratase large subunit |
| VC_RS09960 | VC_2067 | I |  | 2945,5 | 0,42146 | 0,14397 | 2,92745 | 0,003418 | 0,046232 | MinD/ParA family protein |
| VC_RS03820 | VC_0762 | I | *bamB* | 9039,4 | 0,40907 | 0,14098 | 2,90150 | 0,003714 | 0,048790 | outer membrane protein assembly factor BamB |
| VC_RS03255 | VC_0642 | I | *nusA* | 23291,6 | -0,41034 | 0,13618 | -3,01317 | 0,002585 | 0,036913 | transcription termination factor NusA |
| VC_RS12560 | VC_2604 | I | *slyD* | 12166,4 | -0,41416 | 0,13397 | -3,09154 | 0,001991 | 0,031330 | peptidylprolyl isomerase |
| VC_RS16075 | VC_A0558 | II |  | 1932,5 | -0,43790 | 0,14384 | -3,04433 | 0,002332 | 0,034549 | gamma-glutamyltransferase family protein |
| VC_RS03775 | VC_0753 | I | *fdx* | 1523,6 | -0,44634 | 0,15290 | -2,91922 | 0,003509 | 0,047108 | ISC system 2Fe-2S type ferredoxin |
| VC_RS05620 | VC_1146 | I |  | 4127,1 | -0,44872 | 0,15402 | -2,91350 | 0,003574 | 0,047466 | GrxA family glutaredoxin |
| VC_RS09845 | VC_2044 | I |  | 4029,7 | -0,45446 | 0,14711 | -3,08934 | 0,002006 | 0,031330 | Grx4 family monothiol glutaredoxin |
| VC_RS01810 | VC_0359 | I | *rpsL* | 230456,4 | -0,45903 | 0,13083 | -3,50871 | 0,000450 | 0,009472 | 30S ribosomal protein S12 |
| VC_RS14195 | VC_A0177 | II |  | 1767,1 | -0,46627 | 0,14672 | -3,17805 | 0,001483 | 0,024637 | chromosome segregation ATPase |
| VC_RS13580 | VC_A0036 | II | *sstT* | 9991,6 | -0,47902 | 0,15802 | -3,03139 | 0,002434 | 0,035576 | serine/threonine transporter SstT |
| VC_RS12380 | VC_2568 | I |  | 11232,9 | -0,48267 | 0,15503 | -3,11346 | 0,001849 | 0,029642 | FKBP-type peptidyl-prolyl cis-trans isomerase |
| VC_RS01070 | VC_0218 | I | *rpmB* | 181202,7 | -0,48459 | 0,13053 | -3,71260 | 0,000205 | 0,005026 | 50S ribosomal protein L28 |
| VC_RS04085 | VC_0820 | I |  | 2216,4 | -0,48975 | 0,16087 | -3,04447 | 0,002331 | 0,034549 | M66 family metalloprotease |
| VC_RS02205 | VC_0437 | I | *cgtA* | 3794,1 | -0,49193 | 0,14982 | -3,28340 | 0,001026 | 0,018030 | Obg family GTPase CgtA |
| VC_RS09615 | VC_1998 | I | *msrB* | 1384,2 | -0,49766 | 0,16299 | -3,05338 | 0,002263 | 0,034167 | peptide-methionine (R)-S-oxide reductase MsrB |
| VC_RS18225 | VC_A1035 | II |  | 2754,6 | -0,50465 | 0,16325 | -3,09124 | 0,001993 | 0,031330 | DUF3297 family protein |
| VC_RS01575 | VC__t017 | I |  | 12404,7 | -0,50536 | 0,17374 | -2,90880 | 0,003628 | 0,048011 | tRNA-Thr |
| VC_RS04170 | VC_0838 | I | *toxT* | 1415,2 | -0,50881 | 0,16601 | -3,06500 | 0,002177 | 0,033143 | pilus/toxin transcriptional regulator ToxT |
| VC_RS11905 | VC_2472 | I | *ygfZ* | 10897,9 | -0,51335 | 0,16380 | -3,13392 | 0,001725 | 0,028021 | tRNA-modifying protein YgfZ |
| VC_RS09115 | VC_1889 | I |  | 1174,9 | -0,51986 | 0,17971 | -2,89278 | 0,003819 | 0,049627 | GNAT family protein |
| VC_RS05115 | VC_1040 | I | *cobO* | 1257,4 | -0,52057 | 0,16187 | -3,21596 | 0,001300 | 0,022005 | cob(I)yrinic acid a,c-diamide adenosyltransferase |
| VC_RS09755 | VC_2026 | I | *yceD* | 39708,6 | -0,52954 | 0,15381 | -3,44281 | 0,000576 | 0,011203 | 23S rRNA accumulation protein YceD |
| VC_RS03605 | VC_0714 | I |  | 2353,0 | -0,53551 | 0,15512 | -3,45234 | 0,000556 | 0,011019 | DUF1904 family protein |
| VC_RS01855 | VC_0369 | I | *rplI* | 67963,2 | -0,54861 | 0,15942 | -3,44119 | 0,000579 | 0,011211 | 50S ribosomal protein L9 |
| VC_RS01840 | VC_0366 | I | *rpsF* | 124037,8 | -0,55063 | 0,17277 | -3,18710 | 0,001437 | 0,024099 | 30S ribosomal protein S6 |
| VC_RS13675 | VC_A0059 | II |  | 57857,4 | -0,55174 | 0,14487 | -3,80844 | 0,000140 | 0,003715 | Lpp/OprI family alanine-zipper lipoprotein |
| VC_RS01815 | VC_0360 | I | *rpsG* | 71485,8 | -0,55525 | 0,13542 | -4,10027 | 4,127E-05 | 0,001378 | 30S ribosomal protein S7 |
| VC_RS03250 | VC_0641 | I | *rimP* | 7076,3 | -0,55997 | 0,14104 | -3,97021 | 7,181E-05 | 0,002178 | ribosome maturation factor RimP |
| VC_RS10315 | VC_2143 | I |  | 43932,4 | -0,56716 | 0,13759 | -4,12211 | 3,754E-05 | 0,001265 | flagellin |
| VC_RS02010 | VC_0397 | I |  | 4933,8 | -0,57163 | 0,13707 | -4,17023 | 3,043E-05 | 0,001079 | single-stranded DNA-binding protein |
| VC_RS03170 | VC_0627 | I | *erpA* | 630,5 | -0,57857 | 0,17431 | -3,31913 | 0,000903 | 0,016430 | iron-sulfur cluster insertion protein ErpA |
| VC_RS04175 | VC_0839 | I | *tcpJ* | 1600,4 | -0,58046 | 0,16432 | -3,53245 | 0,000412 | 0,008887 | toxin-coregulated pilus pre-pilin peptidase TcpJ |
| VC_RS08850 | VC_1835 | I | *pal* | 47653,9 | -0,58046 | 0,16591 | -3,49855 | 0,000468 | 0,009727 | peptidoglycan-associated lipoprotein Pal |
| VC_RS11610 | NA | I | *rnpB* | 68713,8 | -0,58951 | 0,16173 | -3,64512 | 0,000267 | 0,006357 | RNase P RNA component class A |
| VC_RS04190 | VC_0842 | I |  | 494,9 | -0,59221 | 0,19877 | -2,97944 | 0,002888 | 0,040706 | RDD family protein |
| VC_RS13225 | VC_2746 | I | *glnA* | 48538,2 | -0,59724 | 0,19416 | -3,07602 | 0,002098 | 0,032211 | glutamate--ammonia ligase |
| VC_RS10310 | VC_2142 | I |  | 5491,8 | -0,59982 | 0,13462 | -4,45564 | 8,364E-06 | 0,000376 | flagellin |
| VC_RS02195 | VC_0435 | I | *rplU* | 98498,4 | -0,62268 | 0,13489 | -4,61621 | 3,908E-06 | 0,000191 | 50S ribosomal protein L21 |
| VC_RS12270 | VC_2545 | I | *ppa* | 29207,1 | -0,63109 | 0,17402 | -3,62664 | 0,000287 | 0,006785 | inorganic diphosphatase |
| VC_RS03275 | VC_0646 | I | *rpsO* | 63738,7 | -0,63429 | 0,21437 | -2,95886 | 0,003088 | 0,042724 | 30S ribosomal protein S15 |
| VC_RS03245 | VC__t048 | I |  | 5768,1 | -0,64299 | 0,21594 | -2,97772 | 0,002904 | 0,040706 | tRNA-Met |
| VC_RS05840 | VC_1193 | I |  | 2832,3 | -0,64366 | 0,18212 | -3,53417 | 0,000409 | 0,008887 | hypothetical protein |
| VC_RS10105 | VC_2098 | I |  | 839,1 | -0,65041 | 0,15993 | -4,06692 | 4,764E-05 | 0,001576 | DUF2788 domain-containing protein |
| VC_RS10570 | VC__t081 | I |  | 2259,0 | -0,67930 | 0,20727 | -3,27737 | 0,001048 | 0,018332 | tRNA-Leu |
| VC_RS03600 | VC__t050 | I |  | 1863,3 | -0,68002 | 0,16107 | -4,22190 | 2,423E-05 | 0,000900 | tRNA-Asp |
| VC_RS01850 | VC_0368 | I | *rpsR* | 17479,6 | -0,68374 | 0,21094 | -3,24137 | 0,001190 | 0,020613 | 30S ribosomal protein S18 |
| VC_RS01565 | VC__t015 | I |  | 12619,3 | -0,69078 | 0,22888 | -3,01804 | 0,002544 | 0,036684 | tRNA-Tyr |
| VC_RS08270 | VC_1709 | I |  | 5259,3 | -0,71754 | 0,16526 | -4,34199 | 1,412E-05 | 0,000584 | M16 family metallopeptidase |
| VC_RS09120 | VC_1890 | I |  | 7642,6 | -0,72567 | 0,24458 | -2,96704 | 0,003007 | 0,041923 | NAD(P)/FAD-dependent oxidoreductase |
| VC_RS06330 | VC_1299 | I | *queD* | 3690,3 | -0,73063 | 0,18573 | -3,93377 | 8,362E-05 | 0,002453 | 6-carboxytetrahydropterin synthase QueD |
| VC_RS10700 | VC_2213 | I |  | 92092,1 | -0,73946 | 0,15403 | -4,80076 | 1,581E-06 | 8,715E-05 | OmpA family protein |
| VC_RS01690 | VC__t019 | I |  | 4963,5 | -0,73951 | 0,22617 | -3,26969 | 0,001077 | 0,018746 | tRNA-Gly |
| VC_RS07020 | VC_1450 | I | *rtxC* | 250,9 | -0,74132 | 0,22379 | -3,31264 | 0,000924 | 0,016732 | RTX toxin-activating lysine-acyltransferase RtxC |
| VC_RS10560 | VC__t079 | I |  | 2607,3 | -0,76392 | 0,25041 | -3,05065 | 0,002283 | 0,034196 | tRNA-Leu |
| VC_RS09775 | VC_2031 | I |  | 1978,5 | -0,76772 | 0,16353 | -4,69479 | 2,669E-06 | 0,000141 | SulP family inorganic anion transporter |
| VC_RS17490 | VC_A0874 | II |  | 253,9 | -0,78083 | 0,26982 | -2,89386 | 0,003805 | 0,049627 | hypothetical protein |
| VC_RS01685 | VC__t018 | I |  | 1640,4 | -0,78402 | 0,26760 | -2,92980 | 0,003392 | 0,046056 | tRNA-Gly |
| VC_RS01700 | VC__t021 | I |  | 371,6 | -0,79865 | 0,23368 | -3,41776 | 0,000631 | 0,011905 | tRNA-Met |
| VC_RS00715 | VC_0156 | I | *btuB* | 17311,7 | -0,82272 | 0,15715 | -5,23535 | 1,647E-07 | 1,175E-05 | TonB-dependent vitamin B12 receptor |
| VC_RS03760 | VC_0750 | I | *iscA* | 3757,8 | -0,82339 | 0,18848 | -4,36851 | 1,251E-05 | 0,000536 | iron-sulfur cluster assembly protein IscA |
| VC_RS03750 | VC_0748 | I |  | 30258,7 | -0,83636 | 0,22793 | -3,66927 | 0,000243 | 0,005823 | IscS subfamily cysteine desulfurase |
| VC_RS03735 | VC_0745 | I | *suhB* | 15432,4 | -0,84029 | 0,28807 | -2,91701 | 0,003534 | 0,047108 | inositol-1-monophosphatase |
| VC_RS05255 | VC_1068 | I |  | 353,5 | -0,88562 | 0,25804 | -3,43217 | 0,000599 | 0,011529 | metalloregulator ArsR/SmtB family transcription factor |
| VC_RS03200 | VC_0633 | I | *ompU* | 566755,1 | -0,89302 | 0,21021 | -4,24823 | 2,155E-05 | 0,000828 | porin OmpU |
| VC_RS03755 | VC_0749 | I | *iscU* | 5452,4 | -0,95428 | 0,21881 | -4,36114 | 1,294E-05 | 0,000547 | Fe-S cluster assembly scaffold IscU |
| VC_RS16955 | VC_A0754 | II |  | 1841,9 | -1,03800 | 0,29800 | -3,48318 | 0,000495 | 0,010187 | lipase |
| VC_RS03745 | VC_0747 | I | *iscR* | 6311,6 | -1,04401 | 0,29265 | -3,56739 | 0,000361 | 0,008164 | Fe-S cluster assembly transcriptional regulator IscR |
| VC_RS11500 | VC_2388 | I |  | 727,0 | -1,14423 | 0,17409 | -6,57277 | 4,939E-11 | 6,418E-09 | hypothetical protein |
| VC_RS06470 | VC_1329 | I |  | 1562,4 | -1,16641 | 0,15059 | -7,74580 | 9,498E-15 | 3,142E-12 | outer membrane beta-barrel protein |
| VC_RS10535 | VC_t074 | I |  | 364,4 | -1,35315 | 0,32624 | -4,14768 | 3,359E-05 | 0,001153 | tRNA-Gln |

Legend: IfcSE, standard error of the estimated Fold Change values; stat, statistic values; p-adj, Adjusted p-value.

**Table SII. Sample characterization of the RNA sequencing analysis**

| **Condition** | **Sample** | **Nb reads** | **Nb of reads assigned** | **Mean quality score** | |
| --- | --- | --- | --- | --- | --- |
|  |  |  |  | **Q20 bases** | **Q30 bases** |
| **No PmB** | 1 | 66,500,068 | 28,824,448 | 6.641201 G (99.225182%) | 6.496986 G (97.070474%) |
|  | 2 | 73,951,804 | 32,426,670 | 7.392437 G (99.313873%) | 7.244663 G (97.328594%) |
| **PmB** | 3 | 66,366,754 | 29,089,917 | 6.610773 G (98.901252%) | 6.433118 G (96.243419%) |
|  | 4 | 73,224,628 | 32,047,524 | 7.309020 G (99.283400%) | 7.159279 G (97.249368%) |

**Table SIII. Known antimicrobial resistance genes that are differentially regulated in the presence of subinhibitory concentrations of polymyxin B in *V. cholerae* A1552 in the RNA sequencing analysis**

| **ID** | **Gene** | **Description** | **PmB effect on expression** | **Log_2_(FC)** | **FC** | **Ref.** |
| --- | --- | --- | --- | --- | --- | --- |
| VC_0164 | *vexB* | Multidrug efflux RND transporter permease subunit VexB | up | 0,8187 | 1,764 | (3, 4) |
| VC_0165 | *vexA* | Multidrug efflux RND transporter periplasmic adaptor subunit VexA | up | 0,6443 | 1,563 |  |
| VC_1565 |  | TolC family protein | up | 1,3462 | 2,542 | (5) |
| VC_A0047 |  | HlyD family secretion protein/ Putative multidrug resistance efflux pump (EmrA) | up | 1,0306 | 2,043 | (6) |
| VC_0948 | *rlpA* | Septal ring lytic transglycosylase RlpA | up | 0,4547 | 1,370 | (7) |
| VC_0838 | *tcpN/toxT* | Pilus/toxin transcriptional regulator ToxT | down | -0,5088 | 0,703 | (4) |
| VC_0633 | *ompU* | porin OmpU | down | -0,8930 | 0,538 | (8) |
| VC_2418 | *dsbC* | Bifunctional protein-disulfide isomerase/oxidoreductase DsbC | up | 0,7680 | 1,703 | (9) |
| VC_0034 | *dsbA* | Thiol:disulfide interchange protein DsbA/DsbL | up | 0,8160 | 1,760 | (10) |
| VC_0566 | *vxrB* | Response regulator transcription factor VxrB | up | 0,5584 | 1,473 | (11) |
| VC_0673 |  | Sulfite exporter TauE/SafE family protein | up | 1,0565 | 2,080 | (12) |
| VC_1639 |  | ATP-binding protein, TCS histine kinase | up | 0,8704 | 1,828 | (13) |
| VC_A0020 | *vasX* | Type VI secretion system toxin VasX | up | 0,6522 | 1,572 | (14, 15) |
| VC_1577 | *almG* | Glycine-lipid A transferase AlmG | up | 0,7361 | 1,666 | (16) |
| VC_1578 | *almF* | Lipid A modification system glycine carrier protein AlmF | up | 0,7346 | 1,664 |  |
| VC_1579 | *almE* | Lipid A modification system glycine-protein ligase AlmE | up | 0,7985 | 1,739 |  |
| VC_1319 | *carS* | Histidine kinase CarS | up | 1,6354 | 3,107 | (17) |
| VC_1320 | *carR* | Response regulator transcription factor CarR | up | 1,3072 | 2,475 |  |

Legend: FC, FoldChange; PmB, Polymyxin B; ID, Identification number

11.	Mathieu-Denoncourt A, Duperthuy M. The VxrAB Two-Component System is Important for the Polymyxin B Dependent Activation of the Type VI Secretion System in *Vibrio cholerae* O1 strain A1552. Can J Microbiol. 2023.
